# A direct interaction between the Chd1 CHCT domain and Rtf1 controls Chd1 distribution and nucleosome positioning on active genes

**DOI:** 10.1101/2024.12.06.627179

**Authors:** Sarah A. Tripplehorn, Margaret K. Shirra, Santana M. Lardo, Hannah G. Marvil, Sarah J. Hainer, Karen M. Arndt

**Affiliations:** Department of Biological Sciences, University of Pittsburgh, Pittsburgh, PA USA; UPMC Hillman Cancer Center, University of Pittsburgh, Pittsburgh, PA USA

**Keywords:** Chd1, Rtf1, Paf1 complex, nucleosome remodeling, transcription elongation, chromatin

## Abstract

The nucleosome remodeler Chd1 is required for the re-establishment of nucleosome positioning in the wake of transcription elongation by RNA Polymerase II. Previously, we found that Chd1 occupancy on gene bodies depends on the Rtf1 subunit of the Paf1 complex in yeast. Here, we identify an N-terminal region of Rtf1 and the CHCT domain of Chd1 as sufficient for their interaction and demonstrate that this interaction is direct. Mutations that disrupt the Rtf1-Chd1 interaction result in an accumulation of Chd1 at the 5’ ends of Chd1-occupied genes, increased cryptic transcription, altered nucleosome positioning, and concordant shifts in histone modification profiles. We show that a homologous region within mouse RTF1 interacts with the CHCT domains of mouse CHD1 and CHD2. This work supports a conserved mechanism for coupling Chd1 family proteins to the transcription elongation complex and identifies a cellular function for a domain within Chd1 about which little is known.

## INTRODUCTION

Accurate transcription by RNA Polymerase II (RNA Pol II) requires the regulated alteration of nucleosomes both for access to the underlying DNA sequence and maintenance of chromatin structure. Nucleosomes are repositioned, ejected, or compositionally altered by nucleosome remodeling enzymes in coordination with the transcription cycle and other nuclear events (1,2). First discovered through genetic screens in *S. cerevisiae*, nucleosome remodelers have now been identified in many eukaryotes, and a wealth of information from biochemical and structural studies has provided elegant insights into their modes of catalysis (1,2). Despite these advances, we lack a full understanding of how nucleosome remodelers function *in vivo* and how they are mechanistically coupled to the core molecular processes they ultimately regulate.

Studies in *S. cerevisiae* identified the nucleosome remodeler ATPases Chd1 and Isw1 as important for maintaining proper nucleosome spacing during transcription elongation (3–6). Whereas Isw1 is the ATPase subunit of Isw1a and Isw1b complexes, Chd1 is a monomeric nucleosome remodeler (7,8). Yeast strains deleted of *CHD1* have reduced nucleosome phasing beyond the first nucleosome of the gene body (+1 nucleosome), increased histone turnover at the 3’ ends of genes, altered distributions of transcription-coupled histone modifications such as H3K4me3 and H3K36me3, and elevated levels of aberrant transcripts arising from inappropriate access to the genome by transcription factors (3,5,9–12). Dual loss of Chd1 and Isw1 in yeast exacerbates the effects of single *chd1Δ* and *isw1Δ* mutations and results in severe changes to nucleosome positions across highly transcribed genes, consistent with significant functional overlap (3,5,6). Moreover, both ATPases are required to resolve tightly packed nucleosomes at the 5’ ends of genes (13). For Isw1, these effects are mediated through the Isw1b complex, which has a more prominent effect than Isw1a on nucleosome spacing within gene bodies (3,5,6,13). Chd1 also possesses nucleosome assembly activity that appears to be separable from its remodeling activity and is important for the incorporation of histone variant H3.3 into *Drosophila* chromatin *in vivo* (11,14–16). Consistent with its roles in preserving chromatin architecture during transcription elongation, yeast Chd1 is broadly enriched on the bodies of transcriptionally active genes (17–19).

Among the nine CHD family members in metazoans, CHD1 and CHD2 are most similar to *S. cerevisiae* Chd1 (20). In murine embryonic stem (mES) cells, mCHD2 is enriched at the 5’ ends of highly transcribed genes and shares overlapping peaks with mCHD1 (21). mCHD2 also occupies chromatin marked with H3K36me3 and is distributed across transcribed genes, a feature that is shared with yeast Chd1 and distinguishes it from mCHD1, which is more localized to the 5’ ends of genes in mES cells (22,23). Importantly, mCHD1 and mCHD2 are critical for cellular differentiation and development. mCHD1 regulates pluripotency and self-renewal of mES cells, while mCHD2 is important for neural development (23–26). *CHD1^-/-^*and *CHD2*^-/-^ mice die during development (24,27).

Chd1 employs a twist defect mechanism to induce and then resolve transient DNA bulges in the nucleosome (28–30). Chd1 contains at least five structured regions (Figure 1A): the ChEx region, tandem chromodomains, a two-lobed Snf2-like ATPase domain, a SANT-SLIDE DNA binding domain (DBD), and a helical C-terminal domain (CHCT) (31–34). The ChEx region binds the nucleosome acidic patch and competes with other factors that target this exposed surface (34). The chromodomains distinguish between nucleosomes and free DNA substrates by packing against and inhibiting the ATPase motor (35). The DBD senses extranucleosomal DNA during remodeling in a sequence-independent manner to centrally position nucleosomes on a chromatin template (36–39). The CHCT domain of human CHD1 has been characterized through NMR and has dsDNA and nucleosome binding capabilities *in vitro* (33). However, compared to the other structured domains in Chd1, the CHCT domain is greatly understudied. This domain has been omitted from many biochemical assays, its structure within the context of full-length Chd1 is unresolved, and its functions *in vivo* are unknown (32,33).

**Figure 1.**
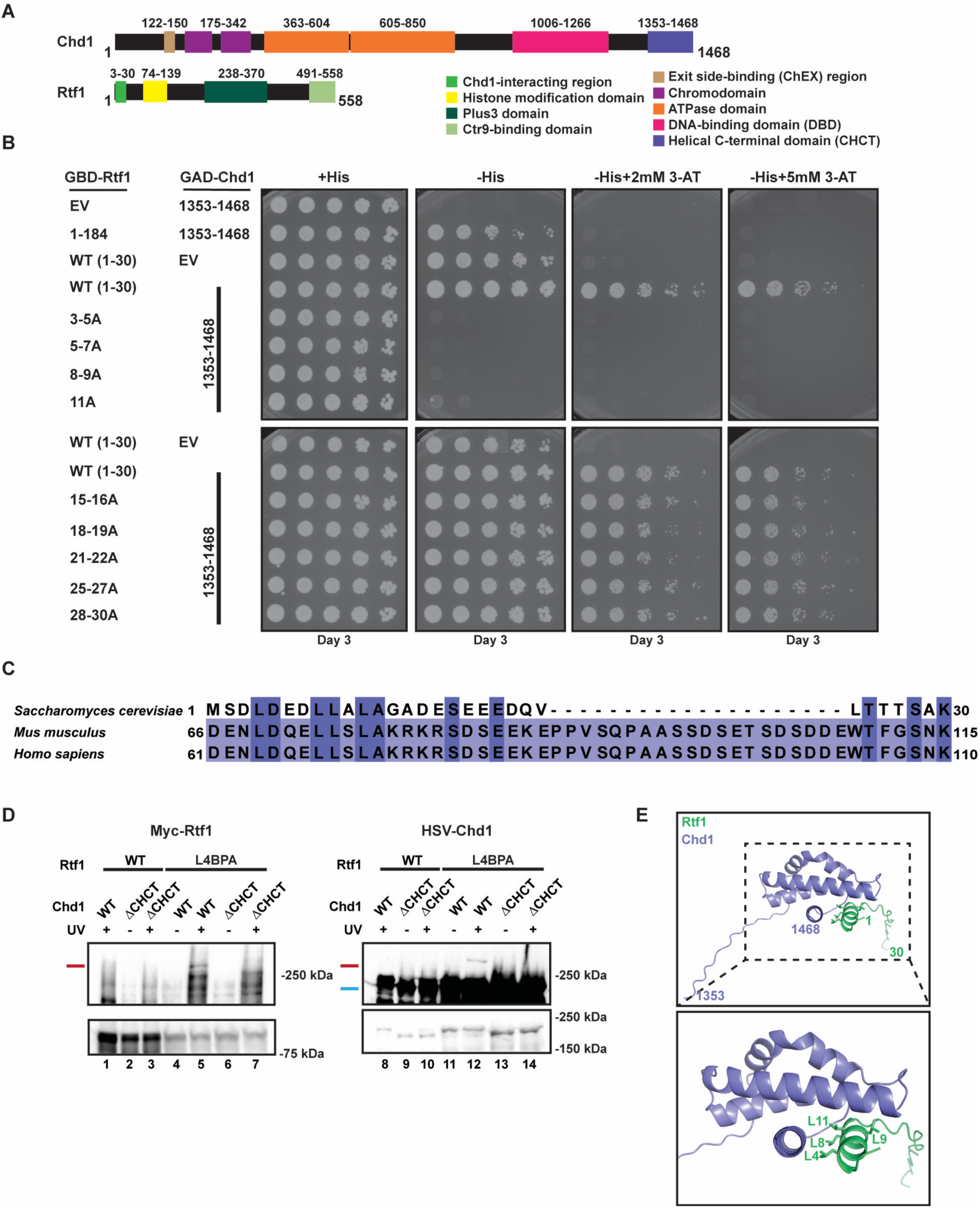
The Rtf1 N-terminal region directly interacts with the Chd1 CHCT domain. (**A**) Diagrams of *S. cerevisiae* Chd1 and Rtf1 with domains indicated (32–34,38,61,72,74,134). (**B**) Y2H analysis of the *S. cerevisiae* Rtf1 N-terminal region and the Chd1 CHCT domain. Plasmids expressing GBD-Rtf1(1-30) with wild-type sequence or the indicated alanine substitutions were co-transformed with a plasmid expressing GAD-Chd1(1353-1468). GBD-Rtf1(1-184), which does not self-activate the *GAL1p-HIS3* reporter, was included as a control. EV = empty GBD or GAD vector as indicated. Cells were plated on indicated media and imaged after three days of growth (n=3). (**C**) Multiple sequence alignment of the *S. cerevisiae* Rtf1 N-terminal region aligned to mouse and human RTF1. (**D**) *In vivo* photo-crosslinking of Rtf1-L4BPA with Chd1. Cells expressing Rtf1 and Chd1 derivatives were exposed to longwave UV light (+) or untreated (-) and then processed for western blot analysis using anti-Rtf1 antisera to detect Myc-tagged Rtf1 and HSV antibody to detect Chd1 derivatives (wild type = WT; C-terminally truncated Chd1 lacking amino acids 1353-1468 = ΔCHCT). The red bars indicate the Rtf1-L4BPA∼Chd1 crosslinked product (lanes 5 and 12). The blue bar indicates Chd1. The panels are from the same membrane that was cut, probed with the indicated antibody, and realigned prior to imaging. Short exposures (bottom) show levels of uncrosslinked Rtf1 and Chd1 in the extracts. (**E**) AlphaFold 3 analysis of the predicted interaction between the Rtf1 N-terminal region (1-30) and the Chd1 CHCT domain (1353 to 1468) (93). Rtf1 amino acids important for the Rtf1-Chd1 interaction by photo-crosslinking or Y2H are indicated.

Elucidation of the mechanisms that target Chd1 to actively transcribed genes is important for understanding how nucleosome spacing and histone modification patterns are maintained during transcription elongation. In addition to their roles in substrate selection, the chromodomains of human CHD1 bind to H3K4me3, a modification that marks the 5’ ends of active genes (40–42). Consistent with this observation, the chromatin occupancy of mouse CHD1 correlates with H3K4me3 and RNA Pol II enrichment at active promoters (22,23). In contrast, the chromodomains of *S. cerevisiae* Chd1 and human CHD2 are unable to bind to H3K4me3 or do so very weakly (42–45), and *Drosophila* CHD1 does not require chromodomains to interact with actively transcribed chromatin (46). These observations suggest the possibility of an alternative, conserved mechanism to direct Chd1 to active genes.

In addition to its nucleosome binding activity, Chd1 interacts with transcription elongation factors and histone chaperones, including Rtf1 and FACT (19,47,48). The ATPase activity of Chd1 is required for the proper distribution of the FACT histone chaperone subunit Spt16 across genes, and Chd1 and FACT coordinately resolve hexasome-nucleosome complexes that arise during transcription elongation (17,49). Rtf1 is a subunit of the Polymerase-Associated Factor 1 Complex (Paf1C), a core component of the active RNA Pol II elongation complex (50,51). Paf1C directly stimulates transcription elongation and RNA Pol II processivity (51–55), facilitates the deposition of transcription-coupled histone modifications including mono-ubiquitylation of H2B on K123 in yeast (K120 in humans) and H3K4me3 (56–62), and impacts RNA splicing and transcription termination (52,63,64). In metazoans, Paf1C regulates promoter-proximal pausing of RNA Pol II (65,66). *S. cerevisiae* Paf1C is composed of five subunits: Paf1, Ctr9, Cdc73, Leo1, and Rtf1; mammalian PAF1C contains a sixth subunit, WDR61/SKI8 (48,50,51,67,68). Paf1C interacts with RNA Pol II through direct contacts with RNA Pol II subunits and the elongation factors Spt5 and Spt6 (51,69–72). In metazoan systems, Rtf1 is more loosely associated with PAF1C (51,73).

An interaction between Chd1 and Rtf1 is intriguing because Rtf1 travels within Paf1C and with RNA Pol II during transcription elongation, providing an avenue by which Chd1 could access and remodel transcriptionally disrupted nucleosomes. We previously demonstrated that Chd1 occupancy at selected genes is reduced when *RTF1* is deleted or mutated in yeast, and a large region of Chd1, that includes the DBD and CHCT domain, can interact with Rtf1 in a manner requiring the Rtf1 N-terminal region (19,74). In *Drosophila*, depletion of Rtf1 reduced Chd1 occupancy on polytene chromosomes, suggesting that an Rtf1-dependent mechanism for targeting Chd1 to active genes could be conserved in other eukaryotes (75). Here, we demonstrate a direct interaction between a 30 amino acid N-terminal region of yeast Rtf1 and the CHCT domain of yeast Chd1. Mutations that disrupt this interaction result in a buildup of Chd1 occupancy at the 5’ ends of genes, a 5’ shift of mononucleosomes and other MNase-protected fragments, and altered histone modification patterns. We further demonstrate an interaction between the analogous regions of mouse RTF1 and mCHD1 or mCHD2 and examine the consequences of disrupting these regions in mES cells. This work provides insight into the mechanisms that distribute the nucleosome remodeler Chd1 across active genes and the function of a conserved Chd1 domain that has been largely uncharacterized.

## MATERIALS AND METHODS

### Yeast strains and media

The *Saccharomyces cerevisiae* strains used in this study are listed in Supplementary Table S1 and are derived from S288C (76) with the exception of PJ69-4A, which was used in the Y2H assays (77). Strains were cultured at 30**°**C in YPD medium supplemented with 400 µM tryptophan or synthetic complete (SC) media lacking indicated amino acids or uracil (78). *Schizosaccharomyces pombe* strains used for spike-in controls were grown in YES medium (3% glucose, 0.5% yeast extract, with adenine, uracil, histidine, leucine, and lysine at 0.2 g/L) at 30**°**C (Supplementary Table S1). For selection against *URA3*, SC medium was prepared with 0.1% 5-fluoroorotic acid (5-FOA) (GoldBio, F-230-10). To select for kanamycin or hygromycin resistance, YPD medium supplemented with 400µM tryptophan and 0.2 g/L Geneticin (Gibco, 11811-023) or 0.3 g/L hygromycin (GoldBio, H-270-1) was used.

Strains containing endogenously expressed *3XHA-rtf1-8-9A*, *3XHA-rtf1-L11A,* and *3xHSV-chd1ΔCHCT* were generated using the *delitto perfetto* approach in which a pCORE cassette that contains a counter-selectable marker is PCR amplified from a plasmid with primers specific to both the cassette and the integration locus (79). To integrate a pCORE cassette at the *RTF1* locus, primers SAT07 and SAT08 containing sequences specific to the pCORE-UH (*K. lactis URA3* and hygromycin resistance) cassette were used. Oligonucleotide sequences are listed in Supplementary Table S2. Stable integration of the pCORE cassettes into a diploid strain generated from crossing KY1691 and KY2822 was confirmed by drug resistance and PCR genotyping. Diploids were then sporulated and haploid strain KY5466 with the pCORE cassette *rtf1Δ::pCORE-UH* was selected for replacement. To integrate a pCORE cassette at *CHD1,* primers containing sequences specific to the pCORE-KU (*K. lactis URA3* and kanamycin resistance) cassette were used. The pCORE cassette was integrated into haploid strain KY3806 to generate *3xHSV-chd1(Δ1353-1468)::pCORE-KU* (KY4361). Stable integration of the pCORE cassette into the strain was confirmed by drug resistance and PCR genotyping. For replacement cassettes, the *RTF1* regions of interest were amplified using primers BTO64 and SAT09 from plasmids KB1653 and KB1654, containing *3xHA-rtf1-8-9A* or *3xHA-rtf1-L11A* in which leucine codons 8 and 9 or 11 were substituted with alanine codons. The *chd1ΔCHCT* truncation was PCR amplified from the *CHD1* locus using SAT79 and SAT81, a primer that encoded a stop codon after amino acid 1352. Amplified fragments were transformed into the appropriate pCORE-containing strain (see above) and recombinants were selected on 5-FOA medium, followed by screening for sensitivity to hygromycin or G418, to generate KY3843, KY3844, and KY4363. Strains were verified by PCR genotyping. Genotypes were further confirmed by inspection of the targeted locus in genomics data. The *GAL1p-FLO8-HIS3* reporter was introduced into strains using genetic crosses as previously described (78).

An HSV epitope-tagged spike-in control strain was generated from *Schizosaccharomyces pombe* strain FWP568 (gift of Fred Winston). A hygromycin resistance cassette (purchased as a plasmid from Addgene *pFA6a-6xGLY-HSV-hphMX4*) (80) was amplified using primers SAT17 and SAT18. Extended homology arms were generated upstream (using primers SAT41 and SAT42) and downstream (using primers SAT43 and SAT44) of the *S. pombe hrp1* locus, which encodes a CHD family protein (81). The homology arms and cassette were assembled using PCR, and the PCR product was used to generate KP08 by integrative transformation of FWP568 (82).

### mES cell culture

ES-E14TG2a (E14) embryonic stem (ES) cells from male *Mus musculus* origin (RRID:CVCL9108) were cultured in Dulbecco’s Modified Eagle Medium (Sigma-Aldrich, D6546-500 mL), 10% Fetal Bovine Serum (Sigma-Aldrich, F2442), nonessential amino acids (Corning, 25-025-CI), 2mM L-glutamine (Corning, 25-005-CI), β-mercaptoethanol, and Leukemia Inhibitory Factor (LIF). Unless otherwise noted, cells were cultured under naïve conditions (serum/LIF +2i), with 3 nM CHIR99021 GSK inhibitor (Med Chem Express, HY-10182/CS-0181) and 1 nM PD0325901 MEK inhibitor (Med Chem Express, HY-10254CS-0062). Cells were passaged every 48 hours using 0.02% trypsin (Corning, 25-052-CI) and quenching with fresh medium. ES cells were grown on plates precoated with 0.2% porcine skin gelatin type A (Sigma Aldrich, G1890) at 37°C in a humified incubator with 5% CO_2_. Routine anti-mycoplasma cleaning was conducted within tissue culture hoods (LookOut DNA Erase spray) and cell lines were screened to confirm no mycoplasma presence.

### CRISPR/Cas9 targeting

mES cell lines were generated with CRISPR/Cas9 technology using single guide RNAs (sgRNAs) and homology constructs that were transfected into low passage wildtype mES cells (Fugene HD, E2311) to produce three cell lines: *chd1ΔCHCT, chd2ΔCHCT,* and *rtf1(AAAA)*. sgRNAs were designed using the CRISPICK algorithm (83,84). sgRNA sequences were purchased as primers and cloned into the px330-U6-Chimeric-BB-CBh-hSpCas9 plasmid containing a PGK-puromycin resistance cassette inserted between NotI and NarI restriction sites (see Supplementary Table S3) (85,86). Homology constructs were purchased from Thermo Fisher Scientific as GeneArt (see Supplementary Table S4 for recombinant homology sequences). Purchased homology constructs were cloned into TOPO vectors using the standard protocol (see Supplementary Table S5 for plasmids used in CRISPR/Cas9 targeting). To generate *chd1ΔCHCT* and *chd2*Δ*CHCT* cell lines, two sgRNAs each were used to generate two cut sites and one homology construct each repaired the DNA breaks. Cell lines were verified by genomic DNA extraction and PCR genotyping using multiplexed primers (see Supplementary Table S3) and checked for appropriate PCR product size. To target *RTF1* and generate the *rtf1*(*AAAA*) cell lines, one sgRNA was used, and clones that had used the homology construct to alter the sequence from LLSL (amino acids 73 to 76) to AAAA were selected. Cell lines were verified by gDNA extraction and PCR genotyping then restriction enzyme digestion with SacI, as the altered DNA sequence resulted in a novel SacI site. Homozygotes for all cell lines were verified by sequencing the PCR products and western blotting to confirm protein expression and relevant protein size shift (Figure 6). Genotypes were further confirmed by inspection of the targeted locus in genomics data. Experiments were performed with two independently generated clones in technical duplicate for *chd1*Δ*CHCT* and three independently generated clones with a single technical replicate for *chd2*Δ*CHCT*, *rtf1*(*AAAA*), and wild type.

### Plasmid construction

To generate plasmids that encode full-length 3xHA-Rtf1 with amino acid substitutions to alanine, plasmid pLS21-5 (87), expressing wild type 3xHA-Rtf1, was digested with EcoRI and BglII or NdeI and BglII (see Supplementary Table S6 for plasmids related to yeast experiments). Primers with substituted sequences (see Supplementary Table S2) were used to amplify from pLS21-5, and fragment inserts were cloned into digested pLS21-5 using Gibson assembly (NEB, E5510S). The plasmid pARL01, a derivative of pGBT9 (88) encoding GBD-Rtf1(1–30), was used as the template to generate plasmids encoding alanine substitutions in the Rtf1 N-terminal region. pARL01 was restriction digested and primers with mutated sequences were used to amplify from pARL01. Plasmids were prepared by Gibson assembly. pGAD-Chd1(863–1468) is from (19). To create pGAD-Chd1(1353–1468), the CHCT region was amplified by PCR from a plasmid containing *CHD1* sequence, using PSO551 and HMO2, and ligated via Gibson assembly into the BamHI and SalI sites in a digested pGAD424 vector backbone (88).

To generate Y2H plasmids with sequences for mouse RTF1, CHD1, and CHD2, DNA fragments encoding CHD1 amino acids 1321-1506, CHD2 amino acids 1331-1561, or RTF1 amino acids 1-321 were codon-optimized for expression in yeast and purchased from Invitrogen (GeneArt; see Supplementary Table S4). Gibson assembly was used to subclone desired fragments into pGBT9 or pGAD424 (88), the GBD-EV or GAD-EV plasmid vectors, respectively, to generate GBD-mRTF1(1–115), GBD-mRTF1(1–321), GBD-mRTF1(66–321), GAD-mCHD1(1321–1506) and GAD-mCHD2(1331-1561). For constructs GAD-mCHD1(1378-1512) and GAD-mCHD2(1432-1567), primers with yeast codon-optimized sequences were used to generate CHD1 and CHD2 fragments by PCR from the GAD-mCHD1(1321-1506) and GAD-mCHD2(1331-1561) plasmids. Plasmid GBD-mRTF11(66-95) was constructed with primers SAT64 and SAT65 that encompass the 66-95 sequence. To construct plasmids encoding alanine-substituted RTF1 sequences, plasmids GBD-RTF1(1-115) and GBD-RTF1(1-321) were used as templates for PCR with primers containing the desired mutations, and the amplified fragments were cloned into linearized pGBT9. See Supplementary Table S2 for the list of primers used to generate codon-optimized Y2H plasmids. All plasmids were verified by DNA sequencing.

### Serial dilution growth assays

Yeast cultures were grown to saturation at 30°C in 5 mL of appropriate media overnight. The optical density (OD) of each culture was measured, and 1 OD unit of cells was diluted in sterile water. Five-fold serial dilutions were prepared and transferred from a sterile 96-well plate to agar media using an 8 x 6 array pinner (Sigma Aldrich, R2383). Plates were incubated at 30°C and imaged daily for at least three days.

### Yeast two-hybrid experiments

Strain PJ69-4A was co-transformed with *TRP1*-marked GBD and *LEU2-*marked GAD plasmids (77). Cultures of transformed cells were grown overnight at 30**°**C in SC-Leu-Trp medium, serially diluted by five-fold, and then plated onto SC-Leu-Trp, SC-Leu-Trp-His, SC-Leu-Trp-His + 2mM 3-aminotriazole (3-AT), or SC-Leu-Trp-His +5 mM 3-AT (3-Amino-1,2,4-triazole, Sigma, 18056-25G). Plates were incubated at 30°C and imaged daily for at least three days.

### Multiple sequence alignments

Protein sequences were retrieved from UniProt, aligned with Clustal Omega using default parameters, and visualized in Jalview (89–91). The following sequences of Rtf1 homologs were used: P53064 (*S. cerevisiae*), A2 (*M. musculus*), and Q92541 (*H. sapiens*). The following sequences of Chd1 homologs were used: P32657 (*S. cerevisiae*), O14139 (*S. pombe*), F4IV99 (*A. thaliana*), O17909 (*C. elegans)*, Q7KU24 (*D. melanogaster*), B6ZLK2 (*G. gallus*), F7D0V6 (*X. tropicalis*), P40201 and E9PZM4 (*M. musculus* CHD1 and CHD2, respectively).

### *In vivo* site-specific photo-crosslinking

BPA (p-benzoyl-L-phenylalanine) crosslinking experiments were performed as described (61,92) except the extracts were made using a urea-based lysis buffer (8M urea, 300 mM NaCl, 50 mM Tris-HCl pH 8.0, 0.1% NP40, 10 mM imidazole, 1 mM PMSF). Yeast strains KY2937 or KY5011 were co-transformed with pLH158/LEU2, a tRNA/tRNA synthetase-expressing plasmid to promote BPA incorporation, and either a plasmid overexpressing wild-type Rtf1 protein (KB851) or a plasmid with an amber codon replacing codon 4 of *RTF1* (KB1424). Yeast cells were grown to late log phase in SC medium lacking leucine and tryptophan and containing 1 mM BPA (BACHEM, F-2800). Twenty OD units of cells were harvested, and crosslinking was performed in 1 mL of ddH_2_O for 10 min with 365 nm UV light. Urea-based lysis buffer (200 µL) was added to the cell pellet and extracts were made by vortexing with glass beads. Lysates were cleared by centrifugation for 15 min in a microcentrifuge and SDS-PAGE loading buffer was added to 1X. Due to the lower expression of Rtf1-L4-BPA, five-fold more extract from these strains was loaded on gels compared to extracts from cells expressing Rtf1. Samples were separated on 4-15% SDS-polyacrylamide gels (Bio-Rad, 4568086), where the 75 kD molecular weight marker was run to the bottom of the gel. Western blot analysis was performed using α-Rtf1 antisera (50) (1:2500) or α-HSV antibody (Sigma-Aldrich, H6030; 1:2000) to detect Chd1 proteins.

### Protein structure prediction

Using standard parameters, AlphaFold 3 was used to predict the structures of Rtf1 N-terminal regions with Chd1 CHCT domains (93). Predictions were performed three times with different seeds. Average IPTM scores (with standard deviation) were: 0.71±0.01 for yRtf1(1–30)-yChd1(1353-1468) and 0.50±0.02 for mRTF1(1-115)-mCHD1(1378-1512). Top scoring models are displayed in the figures.

### Cryptic transcription initiation analysis

Cryptic transcription initiation at the *GAL1p-FLO8-HIS3* reporter was measured as described (9). For experiments involving an *rtf1Δ* strain (KY1370) transformed with *TRP1*-marked *CEN/ARS* plasmids, cells were grown in SC-Trp liquid medium, serial diluted, and spotted onto SC-Trp and SC-His-Trp+gal (galactose, 2%) solid media. For experiments using strains containing integrated *rtf1* or *chd1* mutations, cells were grown in SC complete medium, serially diluted, and spotted onto SC complete and SC-His+gal solid media. Plates were incubated at 30°C and images were taken on the days indicated in the figures.

### ChIP-quantitative PCR (ChIP-qPCR)

*S. cerevisiae* chromatin was prepared as described previously (69). For immunoprecipitation, 700 µL chromatin in 1xFA buffer containing 0.1% SDS, 1 mM PMSF, and 275 mM NaCl was incubated with antibody against HA (40 µL; Santa Cruz HA-probe Antibody (F-7) AC, sc-7392) or HSV (2.5 µL; Sigma-Aldrich, H6030) and incubated overnight at 4°C with end-over-end rotation. Separately, 50 µL chromatin input sample in 1xFA buffer containing 0.1% SDS, 1 mM PMSF, and 275 mM NaCl received no antibody and was incubated overnight at 4°C. The next day, for HSV IPs, 30 µL 50% slurry of Protein A Sepharose beads (Cytiva, 17-5280-01) were washed twice with 1xFA buffer containing 0.1% SDS, 275 mM NaCl, and 1 mM PMSF and added to IP reactions. HSV IPs with Protein A beads were incubated for 1 hr at 4°C with end-over-end rotation. For IP washes, beads were pelleted at 3000 rpm for 10 s and washed for 4 min each in 500 µL of the following buffers: 1) 1xFA buffer containing 0.1% SDS, 275 mM NaCl, and 1 mM PMSF, 2) 1xFA buffer containing 0.1% SDS, 500 mM NaCl, and 1 mM PMSF, 3) TLNNE containing 1 mM PMSF (TLNNE:10 mM Tris-HCl, pH 8.0, 250 mM LiCl, 1 mM EDTA, 0.5% N-P40, 0.5% sodium deoxycholate), and 4) TE buffer (1 mM EDTA, 10 mM Tris-HCl pH 8). IPs were eluted from beads in 50 mM Tris-HCl pH 8, 10 mM EDTA, and 1% SDS at 65°C for 10 min and washed again in TE. Immunoprecipitated and input DNA were treated with Pronase (800 µg/mL) for 1 hr and crosslinks were reversed overnight at 65°C. Immunoprecipitated DNA was then purified using the QIAquick PCR purification kit (Qiagen, 28106) according to manufacturer’s instructions. Final volumes were 100 µL 0.5X TE for IPs and 900 µL 0.5X TE for inputs.

PCR reactions were made in 20 µL volume with 5 µL IP or input DNA, 10 µL SYBR (2x qPCRBIO SyGreen Blue Mix Hi-ROX, 17-505B), 1.2 µL 5 µM primer mix, and water. Reactions were split into 10 µL volumes for technical duplicates. Each sample was tested in biological triplicate and technical duplicate. PCR conditions were: 10 sec initial denature, and 35 cycles of 5 sec 95°C denature and 30 sec 60°C elongation. Primer efficiencies were calculated for each primer pair by performing 10-fold serial dilutions of genomic DNA and performing qPCR in technical triplicate. A line of best fit through the Cq values of the dilution series was obtained and used to calculate primer efficiency. The ChIP-qPCR primers and their efficiencies are found in Supplementary Table S2. Relative chromatin occupancy was calculated as described in (94).

### ChIP-sequencing (ChIP-seq)

Yeast chromatin was prepared as described previously (69). *S. pombe* chromatin from strain KP08 was spiked-in to *S. cerevisiae* chromatin at a 1:9 protein ratio upon thawing of sheared chromatin. Chromatin was quantified using the Pierce BCA kit (Thermo Fisher Scientific, 23225). IPs were performed with 500 µg (for histone IPs) or 1000 µg (for all other IPs) spiked-in chromatin and samples were diluted to 700 µL or 1.4 mL in 1xFA buffer containing 0.1% SDS and 1 mM PMSF, respectively. ChIPs were performed with antibodies against the following epitopes/proteins: HSV (Sigma-Aldrich, H6030; 5 µl per 1000 µg), HA (Santa Cruz, HA-probe Antibody (F-7) AC, sc-7392; 60 µl per 1000 µg), Rpb1 (8WG16; BioLegend, 664906; 4 µL per 1000 µg), H3 (Abcam, ab1791; 1µL per 500 µg), H3K4me3 (Active Motif 39159, 39160; 2.5 µl per 500 µg), and H3K36me3 (Abcam ab9050: 6 µg per 500 µg). After overnight incubation of primary antibodies with chromatin, Protein A Sepharose beads (Cytiva, 17-5280-01) were added to HSV, 8WG16, and H3K4me3 reactions in an amount of 30 µL beads (50% slurry) per 500 µg chromatin. Protein G Sepharose beads (Cytiva, 17-0618-01) were added to H3 and H3K36me3 in an amount of 30 µL beads (50% slurry) per 500 µg chromatin. Samples with Protein A or G beads were rotated end-over-end for 2 hr at 4°C. IP reactions were performed as described above for ChIP-qPCR, with input samples performed in parallel. DNA was eluted in 100 µL 0.5X TE. Libraries were prepared with the NEB Next II Ultra DNA library kit (NEB, E7645L) using 500 pg DNA input. Library quality was verified using fragment analysis (Fragment Analyzer, Advanced Analytical) and submitted for sequencing on an Illumina NextSeq 500 or 2000 instrument for paired-end sequencing to a depth of ∼10 million uniquely mapped reads per sample.

### Indirect immunofluorescence

Yeast cultures were grown to mid-log phase overnight at 30°C in YPD medium supplemented with tryptophan. Cells (5 mL) were centrifuged for 3 min at 3000 rpm and then resuspended in fresh growth medium with 4% formaldehyde, followed by incubation with shaking for 10 min at 30°C. Cells were spun as before and then resuspended in KM solution (40 mM potassium phosphate pH 6.5, 500 µM magnesium chloride) with 4% formaldehyde and then shaken for 1 hr at 30°C. Cells were then spun at 2000 rpm and washed twice in KM solution and once in KMS (40 mM potassium phosphate pH 6.5, 500 µM magnesium chloride, 1.2 M sorbitol). Cells were resuspended in 0.5 mL KMS solution and then treated with 30 µL Zymolyase 100T (US Biological, Z1004250MG) 10 mg/mL stock solution for 25 min at 37°C, followed by centrifugation at 2000 rpm for 2 min. Spheroplasts were washed with KMS solution and then resuspended in 200 µL and placed on ice. Slides were prepared with 1x poly-L-lysine (Sigma Aldrich, P1399) and incubated for 10 min in a humidified chamber, then washed 5 times with water. Twenty µL cells were added to slides and incubated at room temperature for 10 min. Slides were plunged into cold methanol, incubated at -20°C for 6 min, and subsequently plunged into cold acetone for 30 sec. Acetone was evaporated near a heat block at 100°C. Blocking solution (PBS with 0.5% BSA and 0.5% ovalbumin) was applied to the slides for 1 hr at 37°C. After removing the blocking solution with aspiration, a 1:250 dilution of primary antibody (anti-HSV; Sigma-Aldrich, H6030) in PBS (30 µL volume) was placed on the slides overnight at 4°C in a humidified chamber. After removing the primary antibody with aspiration, the cells were washed with blocking solution and incubated with blocking solution for 5 min three times. Alexa Fluor 568 secondary antibody (Invitrogen A-11011; 25 µL) was added to the slides, which were then stored in a dark chamber for 2 hr at 30°C. Slides were washed thrice with blocking solution for 5 min at room temperature. Mounting solution with DAPI stain (Invitrogen, P36966) was applied to the slides and then coverslips were added. Cells were imaged with a confocal microscope (Nikon Confocal, A1R).

### Micrococcal nuclease sequencing (MNase-seq)

The MNase-seq protocol was adapted from (95). Yeast cultures were grown to log phase (OD=0.5-0.6), and cells were crosslinked with 2% formaldehyde, quenched with glycine, pelleted, and flash frozen as described (95). Cell pellets were resuspended in 19 mL sorbitol-Tris buffer (1 M sorbitol, 50 mM Tris-HCl pH 7.4, 125 mM ß-mercaptoethanol) and then spheroplasted by adding 5 mg Zymolyase 100T (US Biological, Z1004250MG) dissolved in 1 mL sorbitol-Tris buffer for 30 min at 30°C while rotating. Spheroplasting efficiency was visually determined under the microscope after mixing 10 µL spheroplasts with 10 µL 10% SDS. Spheroplasts were centrifuged at 4000 rpm for 10 min at 4°C. The supernatant was removed, and the pellet was resuspended to 5 mL in NP buffer (1 M sorbitol, 50 mM NaCl, 10 mM Tris-HCl pH 7.4, 5 mM MgCl_2_, 1 mM CaCl_2_, 0.075% NP40, 1 mM ß-mercaptoethanol, and 500 µM spermidine). MNase (500 units; Sigma, N5386) was resuspended from lyophilized powder in 500 µL Ex50 buffer (10 mM HEPES pH 7.5, 50 mM NaCl, 1.5 mM MgCl_2_, 0.5mM EGTA pH 8.0, 10% glycerol v/v, 1 mM DTT, and 0.2 mM PMSF) and stored in aliquots at -20°C. Spheroplasts in 600 µL aliquots were treated with MNase in a titration ranging from 2 µL to 4.5 µL MNase for 20 min at 37°C with rotation. The reaction was quenched, and samples were treated with proteinase K followed by reversal of crosslinks at 65°C as described (95). DNA was purified by PCI and chloroform isoamyl alcohol extraction followed by isopropanol precipitation. The DNA pellet was resuspended in 40 µL 10 mM Tris-HCl pH 7.5. Resuspended DNA was treated with 5 µL RNase A (Thermo Fisher Scientific, EN0531) for 1 hr at 37°C.

To verify MNase digestion, 15 µL of DNA was purified using the E.Z.N.A. Cycle Pure Kit (Omega Bio-Tek, D6492) and separated on a 2% agarose-TBE gel. The 3.5 µL MNase-treated condition was chosen for all samples based on the titration series, and digested chromatin samples were processed without DNA size selection. *S. pombe* MNase-treated chromatin was separately prepared and mononucleosome-sized DNA fragments were gel extracted and spiked-in to *S. cerevisiae* MNase-digested DNA. Spiked-in MNase-treated DNA was purified using the E.Z.N.A. Cycle Pure kit. Sequencing libraries were prepared with the NEB Next II Ultra DNA library kit (NEB, E7645L) using 50 ng DNA input and 3 PCR cycles. Library quality was verified using fragment analysis (Fragment Analyzer, Advanced Analytical) and submitted for sequencing on an Illumina NextSeq 500 or 2000 instrument for paired-end sequencing to a depth of ∼15 million uniquely mapped reads per sample.

### Western blotting

Yeast protein extracts were prepared either by TCA (for GBD, GAD, and associated G6PDH western blots) (61) or NaOH (96) protein extraction methods (remaining western blots). Proteins were resolved on either 8% or 15% SDS-polyacrylamide gels. Proteins were transferred to nitrocellulose (Bio-Rad, 1620094), with the exception of GBD, GAD, G6PDH, and H3K36me3, which were transferred to PVDF membranes (Immobilon, PVH00010). Membranes were blocked for 1 hr in 5% milk + TBST and subsequently incubated with primary antibodies/antisera against HSV (1:2000), HA (1:5000), Rpb1 (8WG16; 1:500), H3 (1:15,000), H3K4me3 (1:2000), H3K36me3 (1:1000, 3% BSA in PBST), or Sse1 (1:1000) for 2 hr at room temperature in 5% milk + TBST (unless otherwise indicated). Membranes probed with primary antibodies GBD (1:1000), GAD (1:1000), and G6PDH (1:20,000) were incubated overnight at 4°C in 5% milk + TBST.

Western blot analysis of GBD-mRTF1 constructs was performed as follows. Experiments analyzing the expression of yeast Rtf1 fusions to the Gal4 DNA-binding domain (GBD) by western blotting used SC-577 (Santa Cruz). However, this antibody is no longer available, and the tested replacement antibodies reacted non-specifically with proteins that obscured detection of the GBD-mRTF1 proteins on western blots. To circumvent this complication, we performed immunoprecipitations with SC-510-AC (Santa Cruz) on whole cell extracts prepared from strains expressing GBD-mRTF1 and performed western analysis using a different GBD antibody (Sigma-Aldrich, G3042), with the expectation that the cross-reacting background bands of the two antibodies would be different. Cells were grown to OD=0.5-0.6. Extracts were prepared by bead-beating and quantified by Bradford analysis using bovine serum albumin standards. Immunoprecipitations were performed with 500 µg of extracts containing 20 mM HEPES-KOH, pH 7.4, 125 mM potassium acetate, 20% glycerol, 1 mM EDTA pH 8.0, 1 mM DTT, and 1 mM PMSF. Extracts were incubated at 4°C overnight with 15 µL bead volume of SC-510-AC that had been equilibrated with the same buffer. Beads were washed three times with the same buffer and eluted by boiling in SDS-PAGE loading buffer. Approximately one-third of the immunoprecipitation was separated on a 15% SDS-polyacrylamide gel and transferred to a nitrocellulose membrane. Western analysis was performed using the G3042 primary antibody (Sigma-Aldrich) in 5% dry milk in TBST with an overnight incubation at 4°C. To reduce the detection of the heavy and light chain from the immunoprecipitating antibodies, a 1:1000 dilution of Rabbit True Blot (Rockland, 18-8816-13) was used as the secondary antibody. Bands were visualized using chemiluminescence (Thermo Fisher Scientific, 34580) on a Bio-Rad Chemi-doc XRS system.

Mouse protein extracts were prepared by RIPA (86) or acid-based histone extraction. For acid-based histone extraction, cells were lysed in 200 µL Triton Extraction Buffer (TEB; 0.5% Triton X-100, 2 mM PMSF, 0.02% sodium azide diluted in 1x Dulbecco’s PBS) for 10 min. Lysate was centrifuged for 10 min at 650 g at 4°C and supernatant was discarded. Cells were washed in 100 µL TEB and centrifuged as before and supernatant was removed. Lysate was resuspended in 50 µL 0.2 N HCl and rotated overnight at 4°C. Extracts were then centrifuged for 15 min at 4°C and the supernatant was retained, flash frozen, and stored at -75°C. Protein concentrations were determined using the Pierce BCA kit. Extracts were diluted in SDS loading buffer and loaded at either 50 µg protein (CHD1, CHD2, RTF1) or 10 µg protein (H3, H3K4me3, H3K36me3, Actin) onto 8% or 15% SDS-polyacrylamide gels and run at 150V. Proteins were transferred to nitrocellulose membranes. For RTF1 (Invitrogen, PA5-52330), CHD1 (Diagenode, C15410334), and CHD2 (Invitrogen, MA5-47275) blots, loading control was measured with REVERT 700 total protein stain (LICORbio, 926-11011) and imaged by LI-COR (LI-COR Odyssey DLx Imager). Membranes were blocked for 1 hr in 5% milk + PBST. For RTF1 (1:500, 5% milk in PBST), CHD1 (1:2000, 5% BSA in PBST), and CHD2 (1:500 in PBST) blots, membranes were incubated with primary antibody overnight at 4°C. For H3 (1:1000 in PBST), H3K4me3 (1:2500, 5% milk in PBST), H3K36me3 (1:1000, 5% milk in PBST), and actin (1:5000 in PBST) blots, membranes were incubated with primary antibody for 2 hr at room temperature. Following incubation with secondary antibodies (1:5000) for 1 hr at room temperature, membranes were imaged with Pico Plus chemiluminescence substrate (Super Signal West Pico PLUS Chemiluminescent Substrate, Thermo Fisher Scientific, 34580) on the ChemiDoc XRS imaging platform (Bio-Rad).

#### CUT&RUN

CUT&RUN was performed as described using 100,00 nuclei per reaction (86,97–100). Three replicates for wild-type and mutant lines were assayed, except for *chd1*Δ*CHCT* where two technical replicates of two cell lines were used. Nuclei were extracted using a nuclear extraction buffer (20 mM HEPES-KOH, pH 7.9, 10 mM KCl, 0.5 mM spermidine, 0.1% Triton X-100, 20% glycerol, protease inhibitors) and flash frozen. Nuclei were thawed, and nuclei extracted from *Drosophila* S2 cells were added at 1:99 dilution for spike-in normalization. Nuclei were bound to concanavalin A beads (25 uL bead slurry per 100,000 nuclei, Polysciences, 86057-10), blocked with blocking buffer (20 mM HEPES-KOH, pH 7.5, 150 mM NaCl, 0.5 mM spermidine, 0.1% BSA, 2 mM EDTA, protease inhibitors), and washed using a wash buffer (20 mM HEPES-KOH, pH 7.5, 150 mM NaCl, 0.5 mM spermidine, 0.1% BSA, protease inhibitors). Nuclei were incubated overnight at 4°C with rotation in wash buffer with rabbit anti-Chd1 (Diagenode 15410334, 1:50). Nuclei were incubated for 1 hour at room temperature with rotation with rabbit anti-H3K4me3 (Sigma-Aldrich 05-745R, 1:100). Samples were washed twice with wash buffer and incubated for 30 min at room temperature with rotation in wash buffer containing recombinant Protein A-MNase (pA-MN) (1:250). Samples were washed twice with wash buffer and equilibrated at 0°C in an ice bath. Three mM CaCl_2_ was added to each sample to activate MNase cleavage and digested in an ice bath for 30 min. Digestion for H3K4me3 samples was stopped after 30 min using a stop buffer (20 mM EDTA. 4 mM EGTA, 200 mM NaCl, 50 ug/mL RNase A). Samples were incubated at 37°C for 20 min to release genomic fragments and fragments were then separated through centrifugation. Digestion for Chd1 samples was stopped after 30 min using a low salt stop treatment (10 mM EDTA, 2 mM EGTA, 150 nM NaCl, 5 mM Triton-X). Samples were incubated at 4°C for 1 hr to release genomic fragments. The salt concentration was increased to 500 mM for Chd1 samples, RNase A was added, and samples were incubated at 37°C for 20 min to release genomic fragments. Libraries were built using NEB Next II Ultra DNA library kit (NEB E7645L) and AMPure XP beads (Beckman Coulter A63881) purifications. Fragments were PCR amplified with 15 cycles. Samples were then pooled and paired-end sequenced on an Illumina NextSeq2000 to a depth of ∼10 million uniquely mapped reads per sample.

### ATAC-sequencing (ATAC-seq)

ATAC-seq was performed as described (86,101). Nuclei were extracted from 120,000 cells in Lysis buffer (10 mM Tris-HCl pH 7.5, 10 mM NaCl, 3 mM MgCl_2,_ 0.1% NP-40, 0.1% Tween- 20, 0.01% digitonin) Extracted nuclei were flash frozen in liquid nitrogen and stored at -75°C. Nuclei were resuspended in 50 µL Transposase mix (25 µL 2x Tagmentation Buffer, 16.5 µL 1x PBS, 0.5 µL 10% Tween-20, 0.5 µL 1 % Digitonin, 2.5 µL Tagmentase, 5 µL nuclease-free water) and incubated at 37°C for 30 min in a thermomixer shaking at 1000 rpm. DNA was purified and PCR amplified for 9 cycles. Libraries were run on a 1.5% agarose gel and fragments sized 150-700 bp were then gel-extracted. Libraries were paired-end sequenced on an Illumina NextSeq2000 to a depth of ∼50 million uniquely mapped reads per sample.

### ChIP-seq analysis

Integrity of sequencing data was verified by fastqc version 0.11.9. Fastq files were trimmed with trimmomatic version 0.38 with parameters ILLUMINACLIP:TruSeq3-PE.fa:2:30:10 LEADING:3 TRAILING:3 SLIDINGWINDOW:4:15 CROP:36 MINLEN:24 (102). Reads were aligned to a combined genome containing *S. cerevisiae* and *S. pombe* using bowtie2 version 2.4.1 with parameters -q -N 1 -X 1000 for paired-end sequencing. *S. cerevisiae* and *S. pombe* reads were then separated into different bam files for each genome (103). Reads were counted with SAMtools version 1.12 flagstat (104). Normalized bigwigs were generated using deepTools version 3.3.0 bamcoverage with parameters --extendReads --binSize 10 –scaleFactor x, where the scaling factor was determined for each sample using *S. pombe* reads as described (105,106). Normalized bigwigs for two to five biological replicates were averaged using WiggleTools and the ‘mean’ function (107). Pearson correlation values for replicate samples are provided in the Supplementary Data File.

Factor occupancy was evaluated using a bed file spanning the +1 nucleosome dyad to transcription termination site (TTS) with data obtained from (108). Genes that were a minimum of 1 kb in length from the +1 nucleosome dyad to the TTS were analyzed, and we focused on the top quintile of genes with maximum HSV-Chd1 occupancy in our wild-type ChIP-seq data set (n=784 genes). In another size class, we focused on genes that were a maximum of 1 kb from the +1 nucleosome dyad to the TTS and examined the first 500 bp of these genes according to maximum wild-type HSV-Chd1 ChIP-seq occupancy (n=324 genes). To determine regions with maximum factor occupancy for HSV-Chd1 (or for RNA Pol II, as in Figure 2B), the deepTools plotHeatmap function with --sortRegions ‘descend’ and --sortUsing ‘max’ were used to generate sorted bed files (106). Bed files for HSV-Chd1 data were then subset to identify the top quintile of genes and plotted with deepTools computeMatrix and plotHeatmap. To generate metaplots, the output of deepTools plotProfile containing binned data (10 bp) from 200 bp upstream of the +1 nucleosome and 1 kb beyond for each factor and respective genotype was loaded into RStudio version 2022.7.1.554. The maximum data point for each genotype was determined and plotted as a vertical line in GraphPad Prism 10. To generate violin plots, the matrix file from deepTools plotHeatmap containing binned data (10 bp) from 200 bp upstream of the +1 nucleosome and 1 kb beyond was obtained for each factor and respective genotype and was loaded into RStudio version 2022.7.1.554. At each gene for wild type and mutant strains, the point of maximum ChIP-seq enrichment was obtained for each factor. Violin plots with these data were generated in GraphPad Prism 10, where the vertical lines denote median (black line) and interquartile ranges of the data (colored lines).

**Figure 2.**
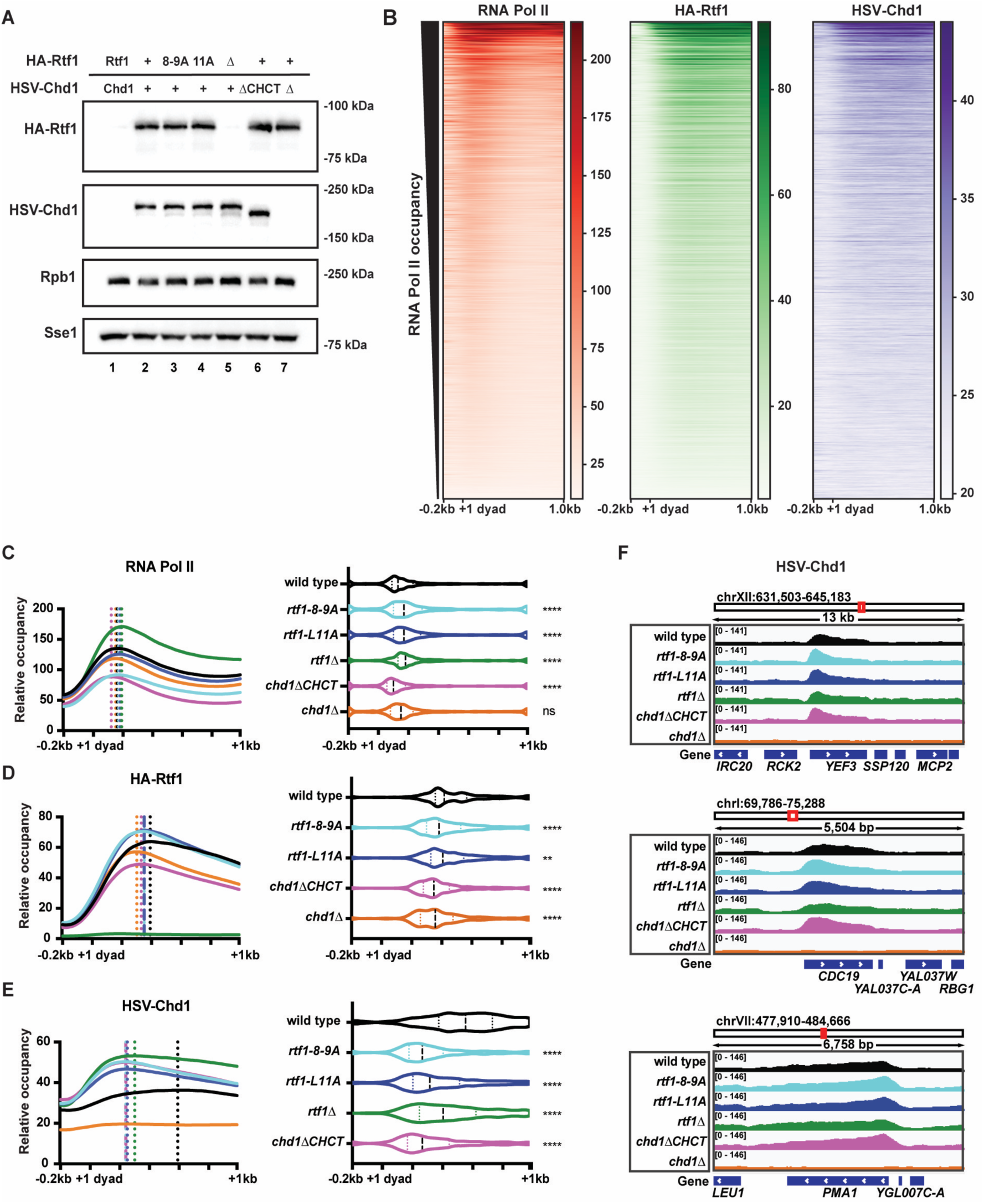
Chd1 distribution across genes is dependent on its interaction with Rtf1. (**A**) Western blot analysis of HA-Rtf1, HSV-Chd1, and Rpb1 levels in strains expressing the indicated HA-Rtf1 and HSV-Chd1 proteins (lanes 2-7; + indicates wild-type tagged protein). In lane 1, extract from an untagged wild-type strain was used as a control. Samples in lanes 5 and 7 were prepared from strains deleted of *RTF1* or *CHD1*, respectively. 8WG16 antibody was used to detect Rpb1. Sse1 served as a loading control. (**B**) Heatmap representations of spike-in normalized RNA Pol II (8WG16), HA-Rtf1, and HSV-Chd1 ChIP-seq data in wild type, sorted by RNA Pol II occupancy for genes over 1 kb in length spanning from 200 bp upstream of the +1 nucleosome dyad to 1 kb downstream (n=3,919 genes). (**C**) Metaplot of average RNA Pol II ChIP-seq signal over the top 20% HSV-Chd1 occupied genes as determined from the wild-type dataset (minimum of 1 kb in length after +1 dyad, n=784 genes). Dotted vertical lines show the point of maximum RNA Pol II occupancy for indicated strains (left). Violin plot representing the distribution of the point of maximum RNA Pol II occupancy for every gene with median and interquartile ranges shown (right). Wilcoxon-rank sum test was used to compare each strain to wild type: p<0.05 (*), p<0.01 (**), p<0.001 (***), and p<0.0001 (****). The average of 2-5 biological replicates for each strain is shown. Information on replicates is provided in the Supplementary Data File. (**D**) As in C, but for HA-Rtf1 ChIP-seq signal (n=784 genes). The average of 2-5 biological replicates for each strain is shown. (**E**) As in C, but for HSV-Chd1 ChIP-seq signal (n=784 genes). The average of 2-4 biological replicates for each strain is shown. (**F**) IGV browser tracks of HSV-Chd1 ChIP-seq signal for genes *YEF3*, *CDC19*, and *PMA1*.

### MNase-seq analysis

Integrity of sequencing data was verified by fastqc version 0.11.9. Fastq files were trimmed with trimmomatic version 0.38 with parameters ILLUMINACLIP:TruSeq3-PE.fa:2:30:10 LEADING:3 TRAILING:3 SLIDINGWINDOW:4:15 CROP:36 MINLEN:24 (102). Reads were aligned to a combined genome containing *S. cerevisiae* and *S. pombe* using bowtie2 version 2.4.1 with parameters -q -N 1 -X 1000 for paired-end sequencing (103). *S. cerevisiae* and *S. pombe* reads were then separated into different bam files for each respective genome. Reads were counted with SAMtools version 1.12 flagstat (104). Due to variability introduced when using *S. pombe* spike-in normalized bigwigs, CPM analysis normalization was used. Normalized bigwigs were generated using deepTools version 3.3.0 bamcoverage with parameters --normalizeUsing CPM --MNase --binSize 1 --smoothLength 10 --minFragmentLength 137 –maxFragmentLength 157 to examine mononucleosome-sized fragments (106). Other MNase-protected fragments were bioinformatically size-selected using bamCoverage, including subnucleosomes (90-110 bp), hexasome-nucleosome complexes (230-270 bp), and dinucleosomes (290-350 bp). Normalized bigwigs for two to five biological replicates were averaged using WiggleTools and the ‘mean’ function (107). Information on replicates is provided in the Supplementary Data File. We generated heatmaps with deepTools computeMatrix and plotHeatmap (106). We analyzed genes that are a minimum of 1 kb in length from +1 nucleosome to TTS and used a bed file that contains only the top 20% of wild-type HSV-Chd1 occupied genes and sorted by maximum HSV-Chd1 occupancy (as described in ChIP-seq data analysis) (n=784 genes). To generate metaplots for each genotype, the matrix obtained from deepTools computeMatrix containing data from 200 bp upstream of the +1 nucleosome, called from (108), and 1 kb beyond in 5 bp bins (240 bins total) was loaded into RStudio version 2022.7.1.554. Data for +1 to +4 nucleosomes were subset into 30 bins (where each bin is 5 bp). The nucleosome center was defined by calling the five-maximum consecutive bins for each 30-bin nucleosome subset (+1 through +4) with the middle bin selected as the center point. Metaplots of these data were generated in GraphPad Prism 10 with the nucleosome centers displayed as vertical lines for wild type (black) or mutant (indicated color).

### CUT&RUN analysis

Fastq files were trimmed to 25 bp using an awk command. Integrity of sequencing data was verified by fastqc version 0.11.9. Reads were aligned to the *Mus musculus* (mm10) and *Drosophila melanogaster* (dm6) genomes using bowtie2 version 2.4.5 with parameters -I 10 -X 1000 -N 1 --very-sensitive for paired end-sequencing (103). Low quality reads were filtered using Picard version 2.5.0 and filtered for mapping quality (MAPQ ≥ 10) using SAMtools version 1.14 (104). Reads were counted with SAMtools version 1.14 flagstat. Datasets were computationally size selected using SAMtools, with 150-500 bp for H3K4me3 and 130-500 bp for CHD1. For H3K4me3, normalized bigwigs were generated using bamcoverage (106) with parameters -- centerReads --binSize 5 –scaleFactor x, where the scaling factor was determined for each sample using *Drosophila* reads (109). For CHD1 CUT&RUN, normalized bigwigs were generated using bamcoverage (106) with parameters --centerReads --binSize 5 --normalizeUsing RPGC -- effectiveGenomeSize 2308125349. Normalized bigwigs for three or four replicates were averaged using WiggleTools and the ‘mean’ function (107). Browser tracks were generated from merged bigwig files using IGV genome browser. Published mES cell CHD1 ChIP-seq data (SRR1747927/SRR1747928) were subset to identify the top quintile of Chd1-bound genes and plotted with deepTools computeMatrix and plotHeatmap. Published mES cell MNase-seq data (SRR1265799) was used to compare profiles, with normalized bigwigs generated using bamcoverage (106) with parameters --centerReads --binSize 5 --normalizeUsing RPGC -- effectiveGenomeSize 2308125349. To generate metaplots, the output of deepTools plotProfile containing binned data (10 bp) from TSS to 2 kb or -1 kb to +2 kb. Output from plotProfile was plotted in GraphPad Prism 10 to generate final metaplots. The mm10 Encode Blacklisted regions were also excluded from the normalized bigwigs. Information on replicates is provided in the Supplementary Data File.

### ATAC-seq analysis

ATAC-seq data quality was verified with Pepatac version 0.10.3 (110). Pepatac parameters were: -Q paired -M 90G --aligner bowtie2 --trimmer trimmomatic --deduplicator picard --peak-caller macs2 --peak-type fixed -G mm10 -P 24 -R (110). De-duplicated bam files obtained from Pepatac were shifted for ATAC-seq analysis using the deepTools alignmentSieve function using alignmentSieve –ATACshift (106). Resulting bam files were then size-selected into accessible fragments (length 1 to 100bp) or mononucleosome-sized fragments (length 180 to 247 bp) using deepTools bamCoverage with settings --normalizeUsing RPGC --effectiveGenomeSize 2652783500 --MNase --binSize 1 –extendReads and by removing the mm10 Encode Blacklisted regions (101). Normalized bigwigs for three to four replicates were averaged using WiggleTools and the ‘mean’ function (107). Published CHD1 ChIP-seq datasets and an mES cell MNase-seq dataset were used as described in the CUT&RUN analysis section. Output from plotProfile was plotted in GraphPad Prism 10 to generate final metaplots. NucleoATAC was used to call nucleosomes in wild type replicates (111). NucleoATAC-called nucleosomes from two or more wild-type replicates were then selected. Nucleosomes over all genes or CHD1 top quintile occupied genes were subset using bedtools intersect and used to assess mononucleosome ATAC-seq enrichment in wild type and mutant cell lines. MNase-seq data are overlayed as a control for nucleosome enrichment at these sites. Information on replicates is provided in the Supplementary Data File.

### Data reproducibility

All sequencing experiments were performed for 2-5 biological replicates with replicate information provided in the Supplementary Data File. Yeast ChIP-seq experiments were spiked-in with *S. pombe* chromatin. H3K4me3 CUT&RUN experiments were spiked-in with batch-controlled *Drosophila* nuclei. For all experiments on mutant strains or cell lines, wild-type samples were grown and processed in parallel to minimize batch effects. Pearson correlation values are reported for all sequencing experiments in the Supplementary Data File. Western blot, indirect immunofluorescence, ChIP-qPCR, and Y2H experiments were performed in biological triplicate. The BPA crosslinking experiment in Figure 1D was performed in biological duplicate. Error bars for ChIP-qPCR represent standard deviation with unpaired t-tests comparing wild-type to mutant strains.

## RESULTS

### An N-terminal region of Rtf1 is sufficient to interact with the Chd1 CHCT domain

We previously found that amino acids 863-1468 of *S. cerevisiae* Chd1 can interact with full-length Rtf1 in a yeast two-hybrid assay (Y2H), but an Rtf1 derivative lacking amino acids 3-30 cannot support this interaction (19,74). To determine the region of Rtf1 sufficient for its interaction with Chd1, we performed a Y2H experiment with the Gal4 Activation Domain (GAD)-Chd1(863–1468) construct, a series of Gal4 DNA Binding Domain (GBD)-Rtf1 constructs, and a yeast strain containing a *GAL1p-HIS3* reporter gene (Supplementary Figure S1A) (77). We observed interaction between GAD-Chd1(863-1468) and full-length GBD-Rtf1, GBD-Rtf1(1-184), and GBD-Rtf1(1-30) as indicated by growth on -His medium or -His medium supplemented with 3-aminotriazole (3-AT) to elevate the stringency of the assay (Supplementary Figure S1A). Empty vector (EV) controls showed that both Rtf1 and Chd1 are required for growth on the test media except for the GBD-Rtf1(1-30) construct, which supported growth on -His medium even in the presence of the GAD empty vector (Supplementary Figure S1A). With 3-AT included in the media, the background signal from self-activation by GBD-Rtf1(1-30) was eliminated, allowing us to observe interactions between GBD-Rtf1(1-30) and GAD-Chd1(863-1468) or additional Chd1 constructs (below).

Chd1(863-1468) contains the Chd1 DBD, an adjacent disordered region, and the structured CHCT domain (Figure 1A). We hypothesized that the CHCT domain may be required for the Rtf1-Chd1 interaction and created a GAD-Chd1(1353-1468) construct. In this Y2H experiment, GAD-Chd1(1353–1468) is sufficient to support an interaction with GBD-Rtf1(1-184) or GBD-Rtf1(1-30) (Figure 1B), demonstrating that the Chd1 CHCT is the domain interacting with Rtf1.

To identify the amino acids in Rtf1 required for its interaction with Chd1, we performed an alanine substitution screen across Rtf1(1-30) and tested for interaction with Chd1(1353-1468) by Y2H (Figure 1B,C). Substitutions 3-5A, 5-7A, 8-9A, and 11A in the Rtf1(1-30) construct greatly reduced the interaction with GAD-Chd1(1353-1468) (Figure 1B), whereas substitutions 15-16A, 18-19A, 21-22A, 25-27A, and 28-30A had little to no effect. Similar results were observed with the Chd1(863-1468) construct (Supplementary Figure S1B). Control experiments with empty vectors confirmed the specificity of the interactions (Supplementary Figure S1C). In addition to eliminating the interaction with Chd1(1353-1468), the 3-5A, 5-7A, 8-9A, and 11A substitutions in Rtf1(1-30) eliminated the self-activation phenotype evident on -His medium lacking 3-AT (Supplementary Figure S1C). Deletion of the *CHD1* gene in the Y2H strain did not eliminate self-activation by Rtf1(1-30); therefore, the source of this phenotype remains unclear (Supplementary Figure S1D). Western blot analysis confirmed expression of GBD-Rtf1(1-30) and the mutant derivatives as well as GAD-Chd1(863-1468) and GAD-Chd1(1353-1468) (Supplementary Figure S1E-G). Together, these data demonstrate an interaction between the Chd1 CHCT domain and amino acids 1-30 of Rtf1.

### Site-specific protein crosslinking demonstrates a direct physical interaction between Rtf1 and Chd1 *in vivo*

Multiple sequence alignment of the Rtf1 N-terminus revealed a patch of conserved amino acids that are shared between yeast, mouse, and human, including yeast residues L4, D5, L8, L9, and L11 that are required for the Rtf1-Chd1 interaction (Figure 1B,C). We termed this region the “LLALA box” after the yeast sequence. Using a system for site-specifically incorporating an unnatural amino acid within a protein of interest through amber codon suppression (112), we replaced Rtf1 L4 with the crosslinking competent, photo-reactive phenylalanine analog BPA to test for a direct interaction with Chd1 *in vivo*. Western blot analysis showed a super-shifted UV-dependent product with Myc-Rtf1 L4BPA that was visible when probing for either Rtf1 or HSV-tagged Chd1, which was expressed from the endogenous locus (Figure 1D, lanes 5 and 12). We generated a strain containing an integrated *HSV-chd1ΔCHCT* mutation by deleting the sequence encoding amino acids 1353-1468 of HSV-Chd1 through homologous recombination. The Rtf1-Chd1 interaction is dependent on the Chd1 CHCT domain, as the ΔCHCT mutant did not yield a photo-crosslinked product with Rtf1 L4BPA (Figure 1D, lanes 7 and 14). Therefore, the Rtf1-Chd1 interaction is direct and requires Rtf1 L4 and the Chd1 CHCT.

We modeled the interaction between the Rtf1 N-terminal region (1-30) and the Chd1 CHCT domain (1353-1468) using AlphaFold 3 (93). The Rtf1 N-terminal region, which is not visible in existing structures of Rtf1 (51,113), is predicted to fold into an alpha helix in which L4, L8, and L11 face the C-terminal helix of the CHCT domain (Figure 1E). Thus, Rtf1 amino acids important for the Rtf1-Chd1 interaction as measured by Y2H and *in vivo* photo-crosslinking are predicted to interface with the CHCT domain (Figure 1E).

### Rtf1 regulates the distribution of Chd1 on active genes

To determine the effects of mutating the Rtf1-Chd1 interface on protein localization and chromatin structure *in vivo*, we generated yeast strains expressing integrated, epitope-tagged derivatives of HSV-Chd1 and HA-Rtf1. We first tested if the tags alter the functions of the wild-type proteins using the *GAL1p-FLO8-HIS3* reporter for cryptic transcription initiation (9). Upon changes to local chromatin structure, an otherwise cryptic transcription initiation site within the *FLO8* sequence can become accessible, resulting in a *FLO8-HIS3* fusion transcript that supports growth on SC-His+gal medium (9,114). While *chd1Δ* and *rtf1Δ* strains harboring the reporter show the expected His^+^ phenotypes (9), the HSV-Chd1 and HA-Rtf1 strains are phenotypically wild type in this assay (Supplementary Figure S2A). Based on the Y2H results (Figure 1B), multiple sequence alignment (Figure 1C), and AlphaFold 3 prediction (Figure 1E), we focused on conserved residues in the LLALA box that are responsible for supporting the Rtf1-Chd1 interaction and generated strains expressing HA-Rtf1(8-9A) and HA-Rtf1(11A) from the endogenous *RTF1* locus. Western blot analysis showed that levels of the HA-Rtf1(8-9A), HA-Rtf1(11A), and HSV-Chd1ΔCHCT proteins are similar to wild-type controls (Figure 2A). To test if deletion of the CHCT domain altered the subcellular localization of Chd1, we performed indirect immunofluorescence. Both HSV-Chd1 and HSV-Chd1ΔCHCT co-localized with DAPI staining, demonstrating that nuclear localization of Chd1 is retained in the HSV-Chd1ΔCHCT strain (Supplementary Figure S2B).

To determine the genome-wide impact of the Rtf1-Chd1 interaction on the chromatin occupancy of Chd1, Rtf1, and RNA Pol II, we performed spike-in normalized ChIP-seq. As expected, wild-type HSV-Chd1 and HA-Rtf1 are enriched at highly transcribed genes, as defined by RNA Pol II occupancy levels (Figure 2B) (18,61). To test if the Rtf1-Chd1 interaction affects the distribution of RNA Pol II, Rtf1, or Chd1, we examined the localization of these proteins in wild-type and HA-Rtf1(8-9A), HA-Rtf1(11A), HSV-Chd1ΔCHCT, *rtf1Δ*, and *chd1Δ* mutant strains, focusing on the top quintile of Chd1-occupied genes. Changes in protein distribution were assessed by calculating the position of maximum protein occupancy (Figure 2C-E). Relative to wild type, the patterns of RNA Pol II and HA-Rtf1 occupancy were only modestly affected in the mutant strains (Figure 2C,D). In contrast, we observed a marked 5’ shift in the position of maximum HSV-Chd1 occupancy in the HA-Rtf1(8-9A), HA-Rtf1(11A), and HSV-Chd1ΔCHCT strains at genes sorted into two different length classes (Figure 2E and Supplementary Figure S2C). Interestingly, a more moderate shift in HSV-Chd1 occupancy was observed in the *rtf1Δ* strain compared to the Rtf1-Chd1 interaction mutants. Browser tracks of HSV-Chd1 chromatin occupancy over *YEF3*, *CDC19*, and *PMA1* exemplify the patterns observed genome-wide (Figure 2F).

We performed ChIP-qPCR to verify the ChIP-seq results and found that the HA-Rtf1(8-9A) and HA-Rtf1(11A) substitutions caused significantly increased HSV-Chd1 occupancy at the 5’ end of *YEF3* and decreased HSV-Chd1 occupancy at the 3’ ends of *YEF3* and *CDC19* (Supplementary Figure S2D). HSV-Chd1ΔCHCT occupancy was significantly reduced at the 3’ ends of *CDC19* and *YEF3* (Supplementary Figure S2D). Further, *rtf1Δ* led to reduced HSV-Chd1 occupancy at both the 5’ and 3’ ends of *CDC19*, consistent with previous results (Supplementary Figure S2D) (74). Similar trends, though modest, were observed at *PMA1* (Supplementary Figure S2D). ChIP-qPCR at the same loci revealed no significant changes in HA-Rtf1 chromatin occupancy, except for a modest reduction in HA-Rtf1(11A) at the 3’ end of *CDC19* (Supplementary Figure S2E). Together, these results demonstrate the importance of the Rtf1 LLALA box and the Chd1 CHCT domain in regulating Chd1 distribution across active genes in yeast.

### Disruption of the Rtf1-Chd1 interaction causes cryptic intragenic transcription

Chd1 is required for the regular spacing of nucleosomes across genes (3,5,6,115,116). Therefore, we hypothesized that disruption of the Rtf1-Chd1 interaction, required for proper Chd1 localization, would result in cryptic transcription initiation and altered nucleosome positioning. We tested the Rtf1 LLALA box mutants for cryptic transcription using the *GAL1p-FLO8-HIS3* reporter. When expressed from plasmids, HA-Rtf1, but not the HA-Rtf1Δ3-30, 3-5A, 5-7A, 8-9A, and 11A derivatives, complemented the cryptic transcription phenotype of an *rtf1Δ* strain (Supplementary Figure S3A). HA-Rtf1 protein levels were similar in these transformed strains (Supplementary Figure S3B). Similarly, the integrated Rtf1-Chd1 interaction mutants, expressing HA-Rtf1(8-9A), HA-Rtf1(11A), and HSV-Chd1ΔCHCT from endogenous loci, exhibited a cryptic transcription initiation phenotype (Figure 3A). Notably, while *chd1Δ* causes the strongest phenotype in this assay, the cryptic initiation phenotypes of the HA-Rtf1(8-9A), HA-Rtf1(11A), and HSV-Chd1ΔCHCT mutants appear more severe than that of the *rtf1Δ* strain (Figure 3A). This observation suggests that complete loss of Rtf1 has a different effect than loss of the Rtf1-Chd1 interaction alone.

**Figure 3.**
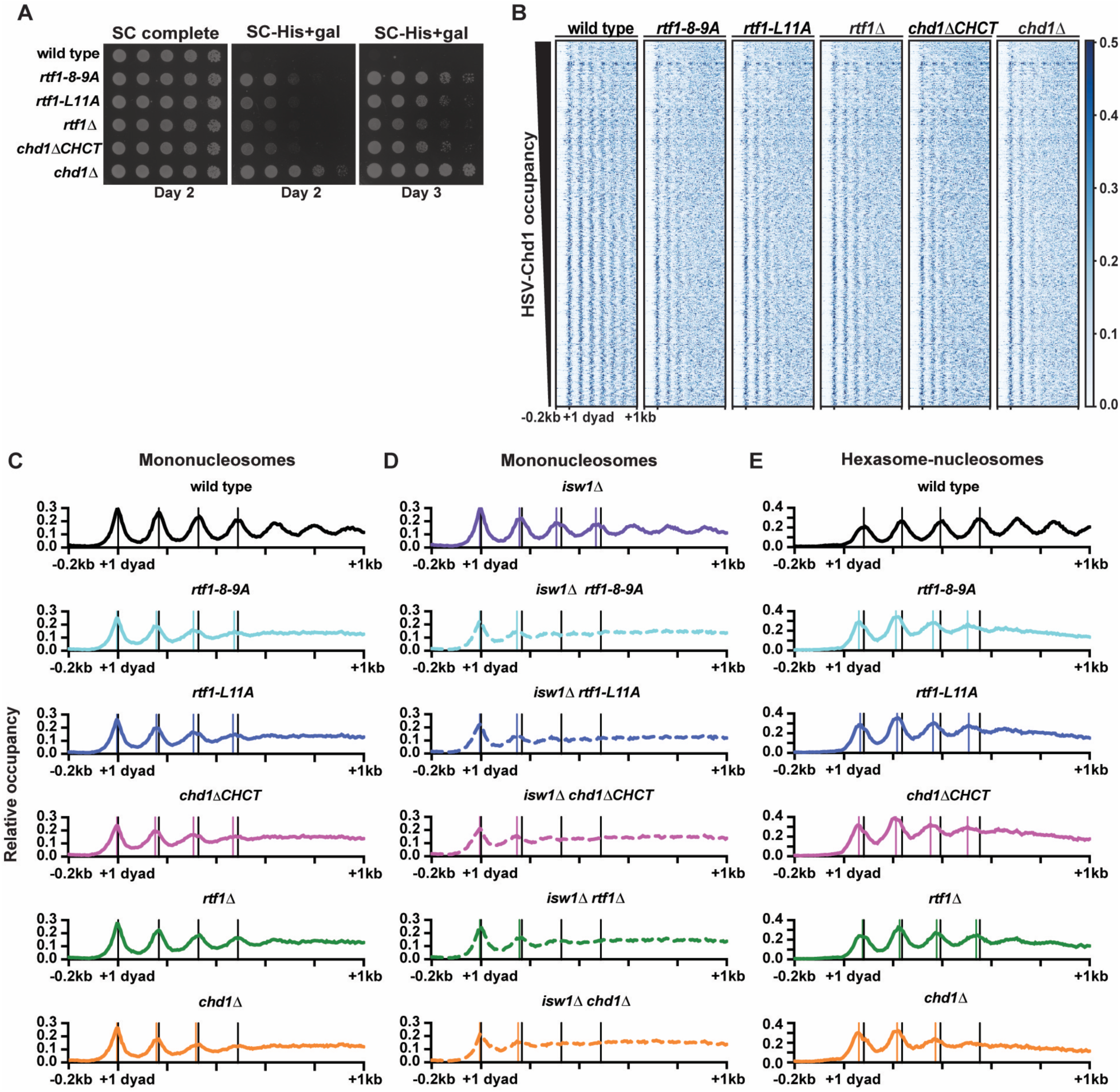
The Rtf1-Chd1 interaction is required for proper nucleosome positioning across Chd1-occupied genes. (**A**) Analysis of cryptic transcription initiation phenotypes of Rtf1-Chd1 interaction mutants using the *GAL1p-FLO8-HIS3* reporter (9). Images were taken on the indicated days after plating (n=3). (**B**) Heatmap representation of mononucleosome positions (137-157bp fragment sizes) determined by MNase-seq over the top 20% of HSV-Chd1-occupied genes as determined from the wild-type dataset (minimum of 1kb in length, n=784). Genes are sorted by HSV-Chd1 occupancy. (**C**) Metaplots of average MNase-seq signal for the genes analyzed in B. Black and colored vertical lines denote the calculated mononucleosome center points of the +1 through +4 nucleosomes in wild-type and mutant strains, respectively. Where no colored lines are shown, no nucleosome shift was observed at that position in mutant strains. Data were divided into 5 bp bins across the region of -0.2 kb to +1 kb relative to the +1 nucleosome dyad, as previously defined (108). Information on replicates is provided in the Supplementary Data File. (**D**) Metaplots of MNase-seq data for mononucleosome positions in *isw1Δ* and indicated double mutant strains as in C. The nucleosome center points for the +3 and +4 nucleosomes in double mutant strains were unable to be calculated. (**E**) Metaplots of hexasome-nucleosome complex (230-270 bp fragment sizes) positions determined from MNase-seq data. Black and colored vertical lines denote the calculated hexasome-nucleosome complex center points in wild-type and mutant strains, respectively. The center point for the +4 hexasome-nucleosome complex in *chd1*Δ was unable to be calculated.

### The Rtf1-Chd1 interaction is required for proper nucleosome positioning over Chd1-occupied genes

To determine if the Rtf1-Chd1 interaction is important for maintaining chromatin structure across genes, we performed MNase-seq on HA-Rtf1(8-9A), HA-Rtf1(11A), HSV-Chd1ΔCHCT, *rtf1Δ*, and *chd1Δ* strains without performing any experimental size selection to enrich for fragments. Relative to wild type, nucleosome positioning is disrupted in all mutants over Chd1-occupied genes (Figure 3B,C). To quantify changes in mononucleosome positions, we calculated the nucleosome center point for the +1 to +4 nucleosomes over the average of Chd1-occupied genes (top quintile, Figure 3C). The nucleosomes in the Rtf1-Chd1 interaction mutants were consistently shifted 5’. Beyond the +4 nucleosome, positioning was severely impacted such that nucleosome locations could not be reliably identified in these strains. In contrast with the Rtf1 point mutants and the Chd1ΔCHCT mutant, the *rtf1Δ* strain showed more modest changes to nucleosome positions, suggesting that substitutions in the Rtf1 LLALA box separate the effect of Chd1 interaction from other Rtf1 functions. These data are in line with the *rtf1Δ* HSV-Chd1 ChIP-seq data, which showed a less pronounced upstream shift of Chd1 (Figure 2E). Similarly, the *chd1Δ* mutant had near wild-type positions at the first three nucleosomes, but a clear loss of nucleosome positioning at and beyond the +4 nucleosome (Figure 3C), consistent with prior studies (3,5,6).

Nucleosome remodeling by Chd1 across gene bodies is partially redundant with nucleosome remodeling by the Isw1 ATPase, as *isw1Δ chd1Δ* double mutants exhibit enhanced changes in genic chromatin structure relative to single mutants (3,5,6,117). Therefore, we hypothesized that the altered nucleosome positioning observed in Rtf1-Chd1 interaction mutants would be exaggerated when paired with *isw1Δ.* At Chd1-occupied genes, *isw1Δ* led to 5’ shifting of nucleosomes, particularly at the +3 and +4 positions (Figure 3D). Deletion of *ISW1* in the HA-Rtf1(8-9A), HA-Rtf1(11A), and HSV-Chd1ΔCHCT strains caused a greater disruption of chromatin structure than observed in the single mutants with 5’ shifts of nucleosomes and near complete loss of nucleosome peaks beyond the +2 position (Figure 3C,D). Nucleosome positioning was also disrupted in the *isw1Δ rtf1Δ* double mutant. These results show that disruption of the Rtf1-Chd1 interaction alters nucleosome positioning across gene bodies likely due to altered Chd1 distribution resulting in improper spacing of nucleosomes.

A recent study demonstrated gene-body enrichment of hexasome-nucleosome complexes (also referred to as overlapping dinucleosomes or OLDNs (118)) and dinucleosomes in yeast strains lacking both Chd1 and Isw1 (49). By focusing on different fragment size classes in our MNase-seq data, we assessed the effects of disrupting the Rtf1-Chd1 interaction on the accumulation and localization of hexasome-nucleosome complexes, dinucleosomes, and subnucleosome particles (Supplementary Figure S3C). Deletion of *CHD1*, in the presence or absence of Isw1, led to a 5’ shift of hexasome-nucleosome complexes and dinucleosomes at the 5’ ends of genes, relative to wild type (Figures 3E and Supplementary Figure S3D, S3E). Disruption of the Rtf1-Chd1 interface by deletion of the Chd1 CHCT domain or substitutions in the Rtf1 LLALA box similarly caused 5’-shifting of hexasome-nucleosome complexes and dinucleosomes to a degree greater than deletion of *RTF1*. In addition, we qualitatively observed an enrichment of the first and second hexasome-nucleosome complexes relative to the third and fourth hexasome-nucleosome complexes in the Rtf1-Chd1 interaction mutants, and this enrichment is not observed in the wild-type strain (Figure 3E). While dinucleosomes and hexasome-nucleosome complexes show a distinct pattern relative to mononucleosomes, subnucleosome positions generally reflect changes observed in mononucleosome positions in the presence or absence of Isw1 (Supplementary Figure S3F). These results demonstrate that the Rtf1-Chd1 interaction is required for the appropriate positioning and resolution of multiple classes of nucleosome particles across gene bodies, and changes to all these particles are enhanced when mutations disrupting the Rtf1-Chd1 interaction are combined with *isw1Δ*.

### Transcription-coupled histone modifications are shifted upstream upon loss of the Rtf1-Chd1 interaction

Histone posttranslational modifications, including H3K4me3 and H3K36me3, are distributed across gene bodies at levels that correlate with transcription (119). Based on our finding that Chd1 occupancy is shifted 5’ with an associated shift in nucleosome positioning in Rtf1-Chd1 interaction mutants, we anticipated a concordant change in transcription-associated histone modifications. ChIP-seq analysis revealed an upstream shift of H3K4me3 and H3K36me3 in the Rtf1-Chd1 interaction mutants and the *chd1Δ* strain relative to wild-type locations at the top 20% of Chd1-occupied genes in two length classes (Figures 4A,B and Supplementary Figure S4A,B). A shift in the broad distribution of total H3 was also observed in the Rtf1-Chd1 interaction and *chd1Δ* mutants, consistent with the MNase-seq data (Figures 4C and Supplementary Figure S4C). In the *rtf1Δ* strain, H3K4me3 was detected at background levels as expected (74), while the H3K36me3 pattern was similar to that of the wild-type strain (Figures 4A,B and Supplementary Figure S4A,B). The shift of H3K4me3, H3K36me3, and H3 in the *chd1Δ* strain agrees with previous results showing that Chd1 is required for their proper distribution (Figures 4A-C and Supplementary Figure S4A-C) (10,12). Browser tracks for *YEF3* and *CDC19* exemplify the shift of H3K4me3 and H3K36me3 in the Rtf1 LLALA box and Chd1ΔCHCT mutants (Figure 4D). Notably, western blot analysis demonstrated wild-type bulk levels of H3, H3K4me3, and H3K36me3 in the HA-Rtf1(8-9A), HA-Rtf1(11A), HSV-Chd1ΔCHCT, and *chd1Δ* mutants (Supplementary Figure S4D). Therefore, mutations that disrupt the Rtf1-Chd1 interaction result in an upstream shift of transcription-coupled histone modifications.

**Figure 4.**
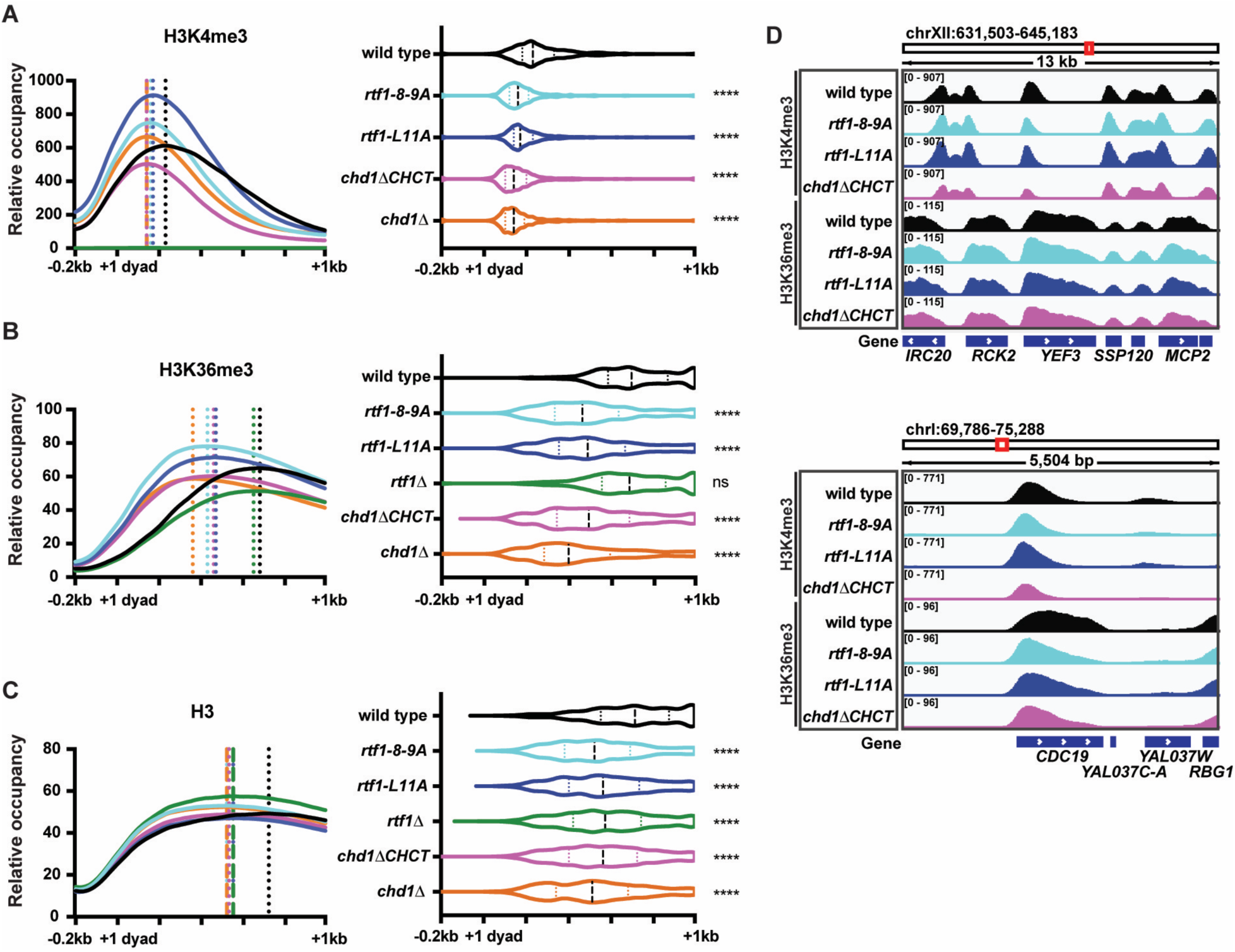
The Rtf1-Chd1 interaction is required for proper localization of modified histones across gene bodies. (**A**) Metaplot of average spike-in normalized H3K4me3 ChIP-seq data from wild-type, *rtf1-8-9A*, *rtf1-L11A*, *rtf1Δ, chd1ΔCHCT*, and *chd1Δ* strains over the top 20% of HSV-Chd1-occupied genes (determined from the wild-type dataset, minimum of 1 kb in length after +1 dyad, n=784 genes) where dotted vertical lines show the maximum point of factor occupancy for each strain (left). The green line represents *rtf1Δ* data. Violin plot representing maximum point of factor occupancy for each individual gene; median and interquartile ranges for the data are shown (right). Wilcoxon-rank sum test was used to compare each strain to wild type: p<0.05(*), p<0.01 (**), p<0.001 (***), and p<0.0001 (****). Data are averaged from 2-5 biological replicates. Information on replicates can be found in the Supplementary Data File. (**B**) As in A, for H3K36me3 ChIP-seq. Data are averaged from 2-3 biological replicates. (**C**) As in A, for H3 ChIP-seq. Data are averaged from 2-3 biological replicates. (**D**) IGV browser tracks of H3K4me3 and H3K36me3 ChIP-seq signals over *YEF3* and *CDC19* in the indicated strains.

### Interaction of mouse RTF1 with CHD1 and CHD2 CHCT domains requires the RTF1 LLSLA box

The direct interaction between the Rtf1 N-terminal region and the Chd1 CHCT domain supports a mechanism by which yeast Rtf1 is required for proper distribution of Chd1 across genes, which in turn leads to appropriate nucleosome positioning and distribution of histone posttranslational modifications. The Rtf1 residues required for this interaction are conserved in mouse and humans (Figure 1C), suggesting that the Rtf1-Chd1 interaction may be conserved. CHD1 and CHD2 are the two mammalian CHD remodelers that are most similar to yeast Chd1, containing conserved domain architectures including CHCT domains (Figures 5A and Supplementary Figure S5A) (33). To test for a potential interaction between RTF1 and CHD1 or CHD2, we conducted Y2H experiments with mouse RTF1, CHD1, and CHD2 proteins using constructs codon-optimized for expression in yeast. Mouse RTF1 contains a sequence similar to the yeast Rtf1 LLALA box, with a sequence change to LLSLA, a 65-amino acid extension N-terminal to the LLSLA box, and an insertion within the region that corresponds to yeast Rtf1(1–30) (Figure 1C). Therefore, we designed several constructs: RTF1(1-321), RTF1(66-321) to eliminate the N-terminal extension, RTF1(1-115) to remove the histone modification domain (HMD) required for H2B ubiquitylation (61,73,74), and RTF1(66–95) which corresponds to yeast Rtf1(1-30) (Figure 5B). For both CHD1 and CHD2, we designed two constructs: the CHCT domain alone and the CHCT domain with an N-terminal extension to the end of the DBD (“extended CHCT”, Figure 5B). As a positive control for the Y2H experiment, we included yeast Rtf1(1-184) and Chd1(1274-1468). EV controls paired with each mouse construct showed no growth on -His medium (Figures 5C and Supplementary Figure S5B). We found that RTF1(1-321) can interact with the CHCT domain alone or the extended CHCT of both CHD1 and CHD2 (Figure 5C). RTF1(66-321) interacted with both CHD1 constructs and the CHD2 extended CHCT construct but did not show an interaction with the CHD2 CHCT domain alone, suggesting that the first 65 amino acids in RTF1 are important for the interactions with CHD1 and CHD2. The smaller RTF1(1-115) construct lacking the HMD was sufficient to interact with all CHD1 and CHD2 CHCT constructs; however, we did not observe interactions between RTF1(66-95) and CHD1 or CHD2. These data demonstrate that the mouse RTF1 N-terminal region can interact with CHD1 and CHD2 via their CHCT domains.

**Figure 5.**
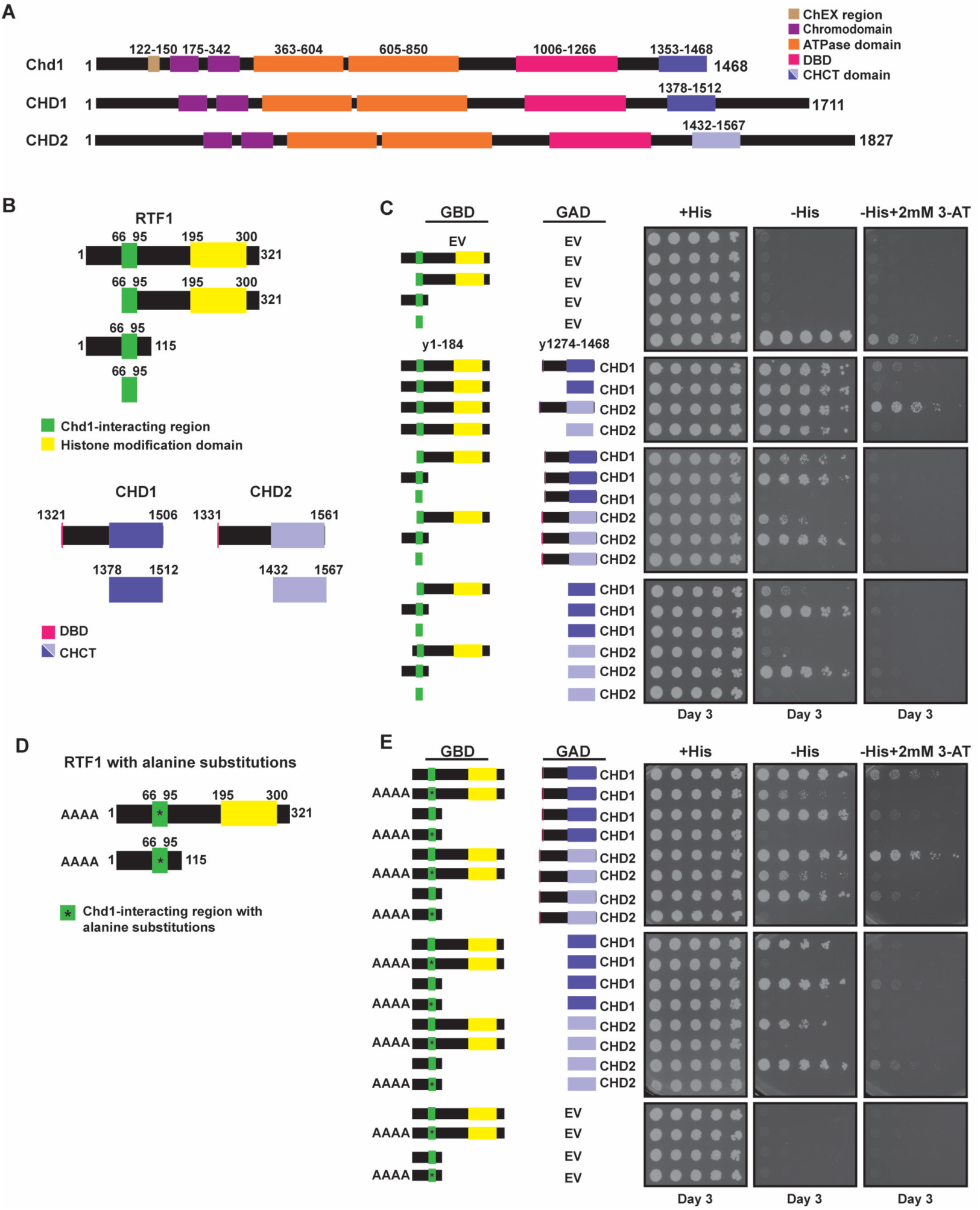
The N-terminus region of mouse RTF1 interacts with the CHCT domains of CHD1 and CHD2 in a LLSLA box-dependent manner. (**A**) Domain organization of *S. cerevisiae* Chd1, mouse CHD1, and mouse CHD2. (**B**) Y2H constructs for mouse proteins RTF1, CHD1, and CHD2. Based on sequence conservation and experimental data in C, the green and yellow boxes indicate positions of the CHD1-interacting region and HMD of mouse RTF1, respectively. (**C**) Y2H analysis of RTF1 with CHD1 or CHD2. Yeast GBD-Rtf1(1–184) and GAD-Chd1(1274–1468) are included as a positive control. (n=3). (**D**) Y2H constructs with alanine substitutions at residues 73-76 within the RTF1 LLSLA box in the context of RTF1(1-321) or RTF1(1-115). (**E**) Y2H analysis of the interactions between wild-type and mutant RTF1 with CHD1 or CHD2 (n=3).

To test if the mouse RTF1-CHD1 and RTF1-CHD2 interactions require the LLSLA box, we introduced alanine substitutions at the LLSL positions in the RTF1(1-321) and RTF1(1-115) constructs (Figure 5D). Demonstrating the importance of the RTF1 LLSLA box for the interactions with CHD1 and CHD2, RTF1(1-115, AAAA) was no longer able to interact with any of the CHD1 or CHD2 CHCT constructs (Figure 5E). We confirmed that levels of the GAD-CHD1 and GAD-CHD2 CHCT proteins were unaffected by the alanine substitutions in RTF1 and that the GBD-RTF1(1-115) and GBD-RTF1(1-115, AAAA) proteins were similarly expressed in the Y2H strains (Supplementary Figure S5C,D). When the LLSLA box was mutated in the longer RTF1(1-321) construct, interactions between RTF1 and the extended CHCTs of CHD1 and CHD2 were partially retained, suggesting that sequences outside the LLSLA box also contribute to these interactions (Figure 5E). Further supporting a conserved interaction between RTF1 and CHD1, we observed an LLSLA-dependent, cross-species Y2H interaction between mouse RTF1(1-115) and the yeast Chd1(863-1468) and Chd1(1353-1468) proteins (Supplementary Figure S5E,F). However, we did not observe a Y2H interaction in the opposite direction, *i.e.* between yeast Rtf1(1-184) and the mouse CHD1 and CHD2 CHCT domain proteins. For RTF1(1-115) and CHD1(1378-1512), AlphaFold 3 predicted a top-scoring model in which the RTF1 helix harboring the LLSLA sequence interacts with the C-terminal helix of the CHD1 CHCT domain (Supplementary Figure S5G), similar to the prediction generated for the yeast Rtf1 and Chd1 proteins (Figure 1E). For RTF1(1-115) and CHD2(1432-1567), an interaction was predicted with only low confidence; however, a recent study focusing on a larger C-terminal fragment of mouse CHD2 reported an RTF1-CHD2 interaction by affinity purification/mass spectrometry supported by an AlphaFold Multimer model (120). Together, these data suggest a conserved interaction between the N-terminal region of RTF1 and the CHD1 and CHD2 CHCT domains.

### mRTF1-CHCT interaction mutants in mES cells minimally affect chromatin structure

Based on our Y2H results supporting a conserved interaction between mouse RTF1 and CHD1/2, we generated mES cell lines to test whether this interaction alters CHD1/2 distribution and chromatin structure across gene bodies. Using CRISPR/Cas9-directed homologous recombination, we generated cell lines encoding either alanine substitutions in the LLSLA box of RTF1, *rtf1(AAAA)*, or deleting the CHCT domains of CHD1 or CHD2 (*chd1ΔCHCT* and *chd2ΔCHCT*). By western blot analysis, we observed minimal changes in RTF1 abundance and the expected changes in the molecular weights of the CHD1ΔCHCT and CHD2ΔCHCT proteins (Figure 6A). Notably, the levels of CHD1 and CHD2 were reduced in the *chd1ΔCHCT* and *chd2ΔCHCT* cell lines, respectively (Figure 6A). To test whether the *chd1ΔCHCT* and *chd2ΔCHCT* mutations alter CHD1 or CHD2 chromatin occupancy, we performed CUT&RUN. While we were unable to obtain high quality CHD2 CUT&RUN datasets after testing three antibodies, we observed enrichment of CHD1 at the 5’ ends of genes in wild-type cells, as previously described using ChIP-seq (Figure 6B and Supplementary Figure S6A,B) (22). In the *chd1ΔCHCT* cell lines, we observed a reduction in CHD1 occupancy likely due to reduced protein abundance. Due to the almost baseline levels of CHD1 in these cell lines, we were unable to robustly determine whether there is a shift in localization at the top quintile of CHD1-occupied genes (as defined from previous ChIP-seq datasets (22)) or over all genes (Figure 6B,C and Supplementary Figure S6A,B).

**Figure 6.**
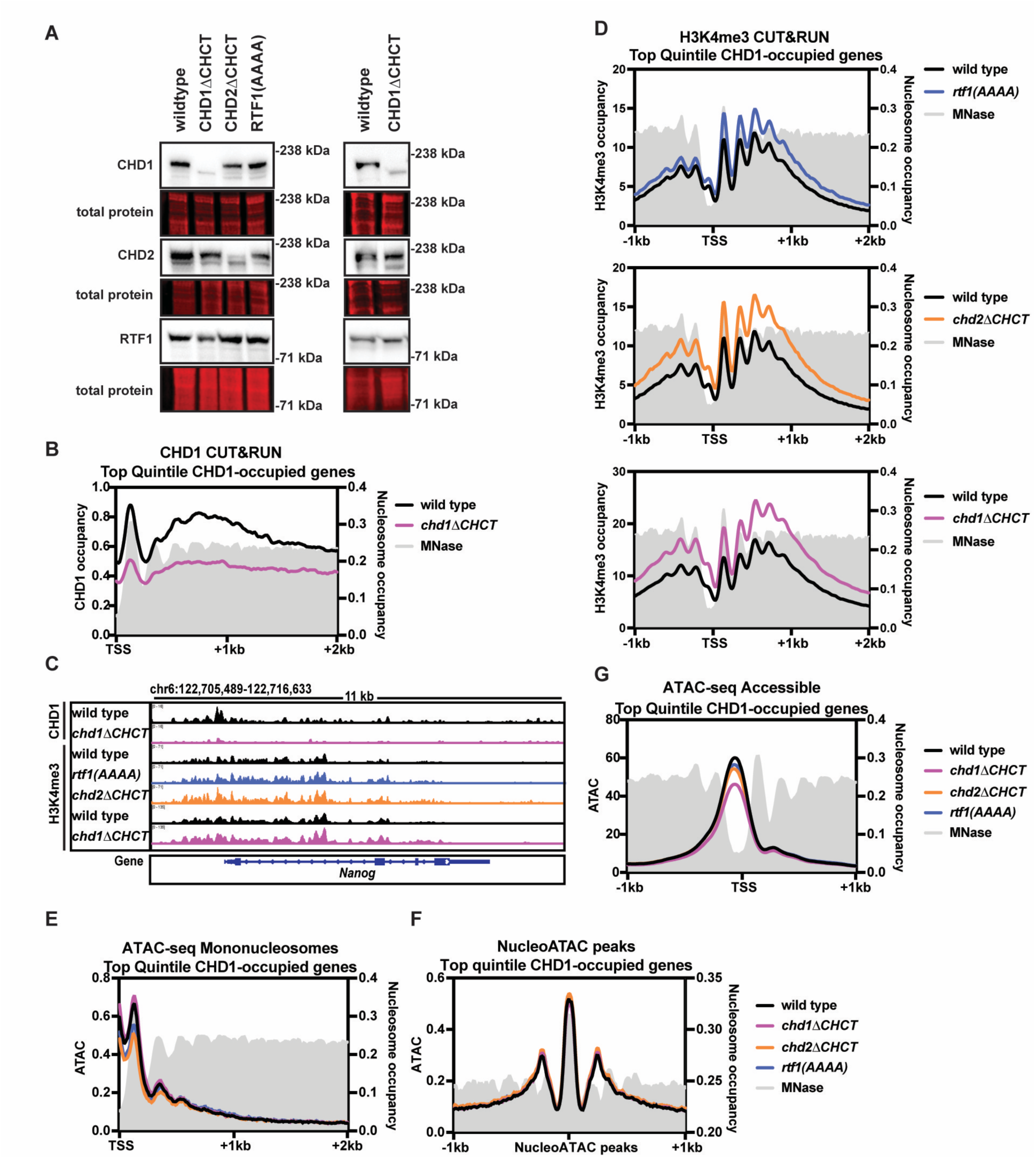
mES cell lines with mutations in the mRTF1-mCHD1/2 interacting regions have minimal changes in H3K4me3 and nucleosome positions. (**A**) Western blot analysis for mouse CHD1, CHD2, and RTF1 in the indicated mES cell lines. Two independent CHD1ΔCHCT clones are shown. (**B**) Metaplot of average CHD1 CUT&RUN data in wild-type (black) and *chd1*Δ*CHCT* (pink) cell lines over the top 20% of CHD1 occupied genes (determined from a wild-type ChIP-seq dataset (22), n=4,101). Data are averaged from 3 wild-type biological replicates and 4 *chd1*Δ*CHCT* replicates (two independent clones assayed in technical duplicate). The gray signal represents mononucleosome data from MNase-seq in mES cells (135). Left y-axis: relative CHD1 CUT&RUN enrichment. Right y-axis: mononucleosome enrichment. (**C**) IGV browser tracks of CHD1 and H3K4me3 CUT&RUN data in indicated cell lines over *Nanog* (left) and *Pou5f1* (right). Data are averaged from 3-4 replicates. Information on replicates can be found in the Supplementary Data File. (**D**) Metaplot of H3K4me3 CUT&RUN in wild-type (black), *rtf1*(*AAAA*) (blue), *chd1*Δ*CHCT* (pink), *chd2*Δ*CHCT* (orange) cell lines as in B. Data are averaged from 3 wild-type, *rtf1*(*AAAA*), and *chd2*Δ*CHCT* biological replicates and 4 *chd1*Δ*CHCT* replicates (as in B). (**E**) Mononucleosome (180-247 bp) ATAC-seq data at genes as in B. Left y-axis: ATAC-seq enrichment. Right y-axis: mononucleosome enrichment as in B. Data are averaged from 3 wild-type, *rtf1*(*AAAA*), and *chd2*Δ*CHCT* biological replicates and 4 *chd1*Δ*CHCT* replicates (as in B). Information on replicates is provided in the Supplementary Data File. (**F**) Mononucleosome (180-247 bp) ATAC-seq data over wild-type nucleosomes from top occupied CHD1 genes called using nucleoATAC (111). The gray signal represents mononucleosome data from MNase-seq in wild-type mES cells. (**G**) ATAC-seq accessible footprints (1-100 bp) as in E.

To test whether nucleosome positions are altered in the *rtf1(AAAA), chd1ΔCHCT*, and *chd2ΔCHCT* cell lines relative to wild type, we performed CUT&RUN for H3K4me3 and ATAC-seq. Total levels of H3, H3K4me3, and H3K36me3 are unchanged in these cell lines (Supplementary Figure S6C). Visual examination of the spike-in normalized H3K4me3 CUT&RUN data demonstrate high resolution nucleosome positions with no visible shift or reduction in occupancy at either the top quintile of CHD1-occupied genes or over all genes (Figure 6C,D and Supplementary Figure S6D,E). The nucleosome-size ATAC-seq reads also show no shift in nucleosome localization in the *rtf1(AAAA), chd1ΔCHCT*, and *chd2ΔCHCT* cell lines relative to wild type, although this may be due to poor nucleosome footprinting beyond the +2 nucleosome (Figure 6E and Supplementary Figure S6F). We then called nucleosome peaks from wild-type ATAC-seq data using nucleoATAC over the top quintile of CHD1-occupied genes or over all genes and assessed the mononucleosome enrichment in all cell lines (Figure 6F and Supplementary Figure S6G) (111). This analysis revealed no global shift in genic nucleosomes in the mutant cell lines. The accessible footprint from these ATAC-seq datasets shows a modest reduction in accessibility over promoters in all three cell lines relative to wild type, with *chd1ΔCHCT* showing the most prominent reduction (Figure 6G and Supplementary Figure S6H). These data suggest that while the interaction between RTF1 and CHD1/2 appears conserved, the chromatin structure over gene bodies in metazoan cells is robust to changes when only a single protein is mutated, likely due to many proteins contributing to nucleosome occupancy and positioning.

## DISCUSSION

In this study, we investigated the mechanistic basis of Chd1 localization to active genes during transcription elongation. We describe a direct interaction between the Paf1C subunit Rtf1 and Chd1, mediated by conserved regions that have been largely uncharacterized. Disruption of this interaction broadly affects Chd1 localization, causes a 5’ directional shift of mononucleosomes and other nucleosome species, and alters histone modification patterns. Our finding that homologous regions in mouse RTF1 and CHD1/2 can interact suggests that mammals and yeast may employ similar mechanisms for coupling nucleosome remodeling to transcription elongation.

Our previous work identified an interaction between Rtf1 and Chd1, demonstrating that Chd1 functions on transcribed genes and is targeted there through interactions with the RNA Pol II elongation complex (19,74). Here, we report a direct interaction between these two proteins and show that a 30-amino acid region of Rtf1 and the CHCT domain of Chd1 are sufficient for this interaction. Despite a wealth of information on Chd1, very little is known about the functions of the CHCT domain. Most biochemical studies on purified Chd1 have omitted this domain, likely for technical reasons, and structural studies have yet to resolve the structure of the CHCT domain in the context of full-length Chd1 or a nucleosome substrate (for example, (32,34,35)). One study on the isolated human CHCT domain revealed a helical bundle fold comprised of five alpha helices by NMR and demonstrated dsDNA and nucleosome binding activity *in vitro* (33). Further, a yeast Chd1 construct lacking the CHCT domain remodels nucleosomes approximately 1.5 times faster than full length Chd1 *in vitro*, suggesting an autoinhibitory role for the CHCT domain (33). Interestingly, some of the conserved residues required for dsDNA binding by the human CHCT domain lie on alpha helix 5, which is predicted in yChd1 and mCHD1 to interface with yRtf1/mRTF1 (Figure 1E and Supplementary Figure S5G). Our data demonstrate the CHCT domain is important for appropriate localization of Chd1 across gene bodies. Given the positioning of Rtf1 within the RNA Pol II elongation complex, it is possible that Rtf1 and DNA compete for binding to the CHCT domain and engage in a handoff mechanism through which nucleosome remodeling is temporally coordinated with transcription elongation. It will be interesting to determine how the dsDNA-binding and Rtf1-interacting functions of the CHCT domain contribute to overall Chd1 function.

The altered distribution of Chd1 in Rtf1-Chd1 interaction mutants is reflected in a 5’ shift in the distribution of different nucleosome species, as defined by the sizes of DNA fragments protected from MNase digestion. Mononucleosome positions in Rtf1 N-terminal and Chd1ΔCHCT mutants shift similarly at the +2, +3, and +4 nucleosomes. Beyond the +4 nucleosome, positioning is severely impacted such that nucleosome positions could not be reliably identified. The more modest change in mononucleosome positions in the *chd1*Δ strain agrees with results from previous MNase-seq studies (3,5). Reflecting the nucleosome shifts quantified using MNase-seq, genic H3K4me3 and H3K36me3 distributions determined by ChIP-seq are also impacted in Rtf1-Chd1 interaction mutants. Consistent with Chd1 and Isw1 having overlapping functions, we observed enhanced nucleosome positioning defects upon introducing the *isw1Δ* mutation into Rtf1-Chd1 interaction mutants. Together, these data reflect the importance of Chd1 in properly spacing nucleosomes such that when Chd1 localization is disrupted, nucleosome positions are shifted in the 5’ direction across gene bodies.

Changes in subnucleosome positions mirror the changes observed with mononucleosomes in the Rtf1-Chd1 interaction mutants. These subnucleosomes may be hexasomes, although further studies are necessary to define their precise composition. Hexasomes lacking distal H2A/H2B dimers are produced at the +1 position in mammals as a consequence of transcription elongation (121), and Chd1 is able to slide hexasomes when the H2A/H2B dimer is on the entry side of the substrate (122). Interestingly, similar to our results with *chd1* mutants, deletion of *INO80*, which encodes the ATPase subunit of the Ino80 nucleosome remodeler complex, also results in a 5’ shift in the positions of mononucleosomes and subnucleosome particles (123).

The hexasome-nucleosome complex consists of a nucleosome that directly abuts a hexasome and has been increasingly studied in recent years (49,118,124,125). Chd1 binds hexasome-nucleosome complexes and, following the FACT-mediated deposition of an H2A/H2B dimer to restore the nucleosome, remodels the new nucleosome to increase the distance between the two nucleosomes (49). We observe a 5’ shift of hexasome-nucleosome complexes in our Rtf1-Chd1 interaction mutants. Dinucleosome-sized fragments, corresponding to tightly packed nucleosomes, were previously reported to accumulate in *chd1Δ isw1Δ* double mutants (13). Our data show that disruption of the Rtf1-Chd1 interaction is sufficient to increase the relative levels of dinucleosomes, especially at the first and second positions, and shift their positioning 5’. Together with our analysis of mononucleosomes and subnucleosome particles, these changes in hexasome-nucleosome complexes and dinucleosomes indicate a broad role for Chd1 in resolving nucleosome species across actively transcribed genes. These molecular changes are reflected in the role of Chd1 in suppressing cryptic transcription (9).

The Rtf1-Chd1 interaction mutants exhibit stronger phenotypes than an *rtf1*Δ strain in several of our assays. The positions of mononucleosomes and other nucleosome species are less severely affected by the full deletion of *RTF1* than by specific mutations that disrupt the Rtf1-Chd1 interaction, although we observe poorly phased nucleosomes beyond the +4 position in both cases. ChIP-seq analysis also revealed a less pronounced shift in H3K36me3 distribution in the *rtf1Δ* strain compared to the *rtf1* point mutants. These results highlight the importance of using specific point and domain mutations to understand the roles of proteins that have multiple functions and interacting partners. The *rtf1Δ* strain reflects the combined outcome of eliminating all Rtf1 functions, including interacting with Chd1, coupling Paf1C to Spt5, and promoting H2B ubiquitylation and downstream modifications such as H3K4me3 (59,61,62,70,72,126,127). It is likely that these functions act coordinately or even in opposition. Therefore, we more specifically capture the consequences of the Rtf1-Chd1 interaction using separation-of-function mutations.

The essential histone chaperone, FACT, which is composed of Spt16 and Pob3 in yeast, is distributed across gene bodies in a manner dependent on Chd1 ATPase activity (17). FACT binds to partially unwrapped H2A/H2B dimers and functions in a transcription-dependent manner (113,128,129). FACT, Chd1, and Paf1C interact in yeast and mammals (19,47,48,130). While modest, Spt16 accumulates at the 5’ ends of genes in *rtf1Δ* cells, suggesting that the 5’ accumulation of Chd1 we observe in *rtf1Δ* may subsequently impact FACT distribution (17).

Through Y2H assays, we found that the interaction between mRTF1-mCHD1 and mRTF1-mCHD2 is conserved and occurs in a mRTF1 LLSLA box-dependent manner. Structure predictions indicate that several conserved leucine residues in yeast Rtf1, including the L4 position mutated for *in vivo* crosslinking and the L8 and L11 positions mutated for our functional studies, face a helix in the Chd1 CHCT. Mutation of analogous mRTF1 residues or deletion of the mCHD1/2 CHCT domain resulted in a loss of interaction by Y2H. Therefore, we hypothesized that mRTF1 LLSLA mutations and mCHD1/2 CHCT deletions would disrupt the predicted interaction in mES cells and shift nucleosome positioning and H3K4me3, as observed in yeast. We were surprised to observe that H3K4me3 localization and mononucleosome positions as assessed by ATAC-seq remained unchanged in RTF1(AAAA), CHD1ΔCHCT, and CHD2ΔCHCT cell lines relative to wild type. Given the large number of nucleosome remodelers in mammals, the mRTF1-CHCT interaction mutations may be insufficient to yield effects at the chromatin level. It may also be that the mRTF1-mCHD1/2 interactions involve sequences outside of the minimally interacting regions such that our specific mutations are insufficient to disrupt these interactions. This possibility is supported by our Y2H results using longer constructs (Figure 5E). Alternatively, for mCHD1, the chromodomain-H3K4me3 interaction may be a dominant driver of CHD1 targeting with the RTF1-CHD1 interaction playing a modulatory role (41,42).

Previous studies using prostate cancer cell lines depleted of hCHD1 or mouse embryonic fibroblasts overexpressing a dominant negative version of mCHD1 reported a reduction in H3K4me3 occupancy and nucleosome occupancy, respectively (131,132). However, we did not observe a reduction in H3K4me3, and in fact observed a modest increase in spike-in normalized H3K4me3 CUT&RUN experiments, despite reduced abundance of mCHD1. These differences may be cell line specific, due to different protein levels in the knockdown versus CHD1ΔCHCT mutant conditions, and/or reflect the need for spike-in normalization. Although we did not observe reductions in nucleosome positioning or H3K4me3 in our mutant mES cell lines, it remains possible that the mRTF1-mCHD1 and mRTF1-mCHD2 interactions detected here support a mechanism for CHD1 recruitment in mammals that is not entirely through H3K4me3 recognition and a potential pathway through which mCHD2, which has weak affinity for H3K4me3, is targeted to gene bodies (43).

In summary, our study shows that a direct interaction between Rtf1 and Chd1 is required for the distribution of Chd1 across transcribed genes in yeast and, as a consequence, the proper distribution of nucleosomes and associated histone modifications. These results demonstrate the necessity for Chd1 and its coupling to the RNA Pol II elongation complex in the re-establishment of chromatin structure in the wake of RNA Pol II. Specifically, our data argue that a functional Chd1-Rtf1 interaction is required to shift all nucleosome species in the 3’ direction, counteracting the retrograde movement of nucleosomes caused by RNA Pol II passage (133). Further characterizing the conservation of this interaction as well as investigating other factors important in this process may yield further insights into the necessity of the RTF1-CHCT interaction in mammalian cells.

## ACKNOWLEDGMENTS

We are grateful to Alex Lederer, Sanhita Kutumbaka, and Matthew Hurton for technical assistance, Craig Kaplan and Fred Winston for providing yeast strains and plasmids, Jeff Brodsky for Sse1 antisera, Greg Bowman for communicating updated information on Chd1 domain boundaries, and members of the Arndt lab, especially Alex Francette, Aakash Grover, and Mitch Ellison, and the Hainer lab for helpful discussions. This research was supported in part by the University of Pittsburgh Center for Research Computing, RRID:SCR_022735, through the resources provided. Specifically, this work used the HTC cluster, which is supported by NIH award number S10OD028483. This project used the University of Pittsburgh HSCRF Genomics Research Core, RRID: SCR_018301 NGS sequencing services, with special thanks to the Assistant Director, Will MacDonald.

