## Supplemental Data and Tables for "A direct interaction between the Chd1 CHCT domain and Rtf1 controls Chd1 distribution and nucleosome positioning on active genes"

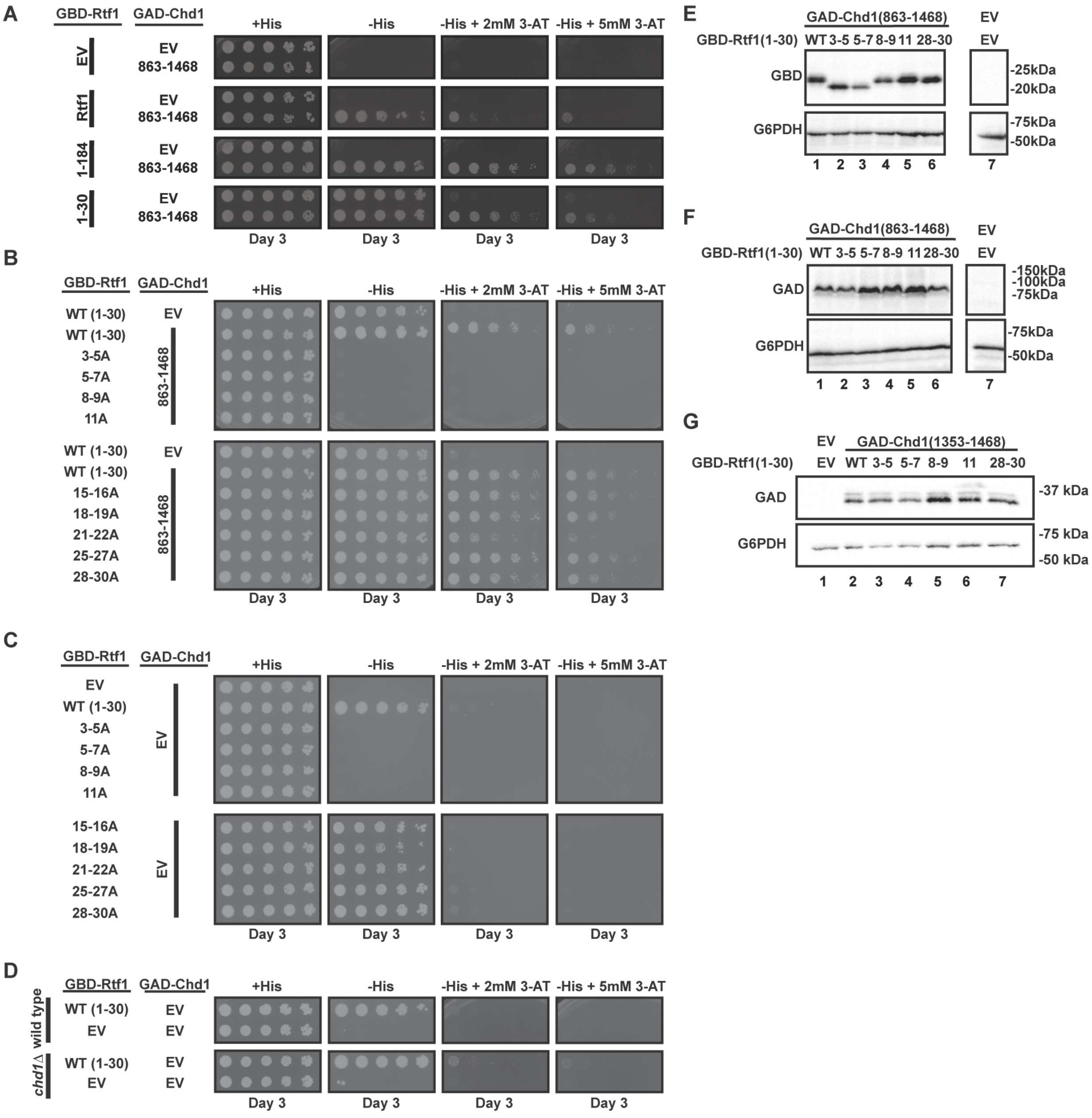


**Supplementary Figure S1. The N-terminal region of Rtf1 interacts with a Chd1 construct containing both the DNA-binding and CHCT domains.** (**A**) Yeast two-hybrid (Y2H) analysis of full-length GBD-Rtf1, GBD-Rtf1(1-184), or GBD-Rtf1(1-30) in combination with GAD-Chd1(863-1468) (n=3) (1). (**B**) Y2H analysis of GAD-Chd1(863-1468) with wild-type GBD-Rtf1(1-30) or GBD-Rtf1(1-30) derivatives containing the indicated alanine substitutions (n=3). (**C**) Y2H analysis of wild-type GBD-Rtf1(1-30) or substitution mutants with the GAD empty vector (EV). (**D**) Y2H analysis of the GBD-Rtf1(1-30) self-activation phenotype in wild-type and *chd1Δ* strains (n=3). (**E**) Western blot analysis using anti-GBD antibody to assess levels of the GBD-Rtf1(1-30) wild-type and mutant proteins in Y2H transformants containing GAD-Chd1(863-1468). Cropped lanes are from the same membrane and are organized for clarity. (**F**) Western blot analysis using anti-GAD antibody to assess the levels of GAD-Chd1(863-1468) in Y2H transformants containing wild-type or mutant GBD-Rtf1(1-30). Cropped lanes are from the same membrane and are organized for clarity. (**G**) Western blot analysis using anti-GAD antibody to assess the levels of GAD-Chd1(1353-1468) in Y2H transformants containing wild-type or mutant GBD-Rtf1(1-30). G6PDH serves as a loading control. EV = empty GBD or GAD vector as indicated.

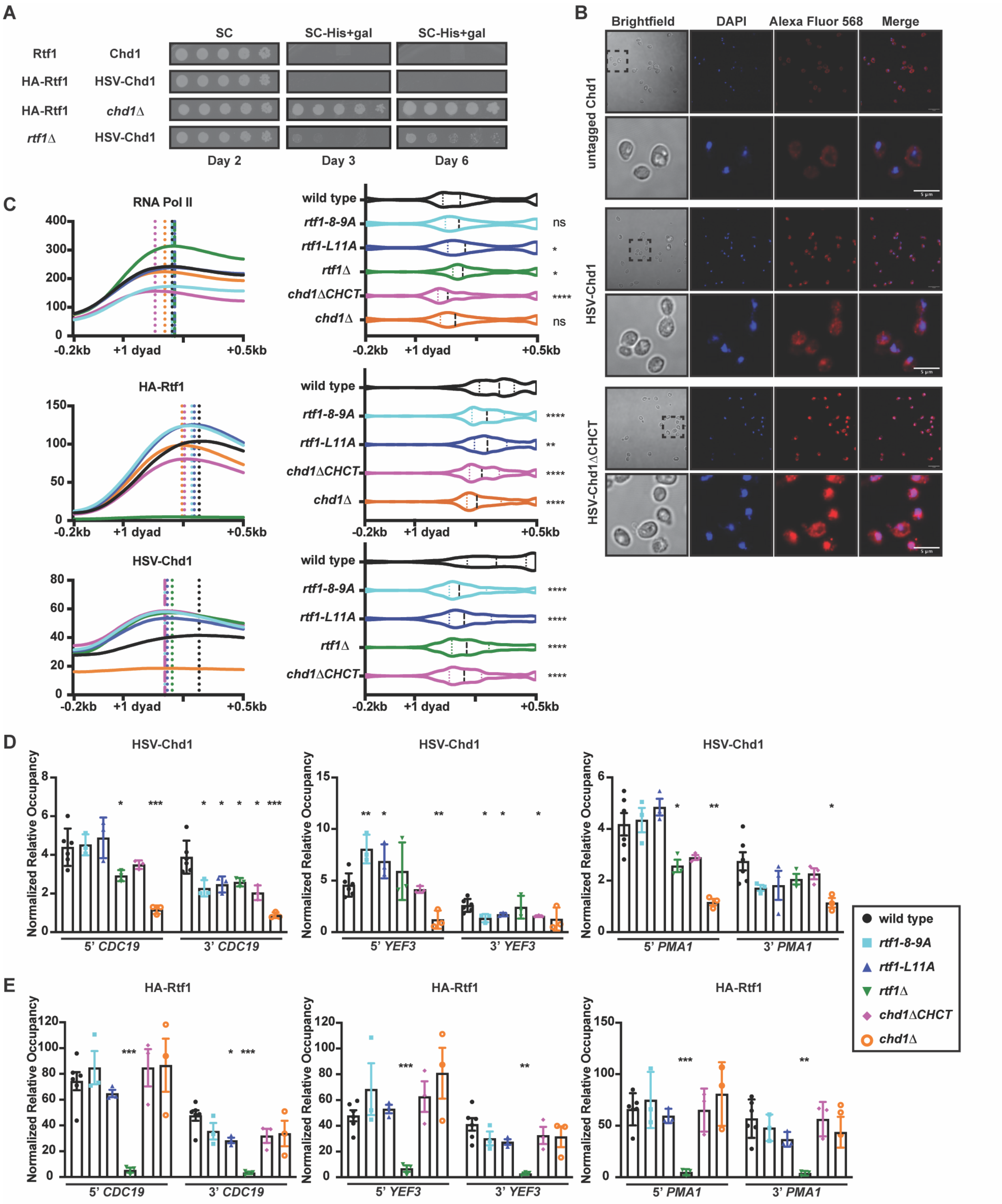


**Supplementary Figure S2. Chd1ΔCHCT localizes to the nucleus and has altered chromatin distribution.** (**A**) Comparison of cryptic transcription initiation phenotypes using the *GAL1p-FLO8-HIS3* reporter (2) for strains expressing untagged or epitope-tagged Rtf1 and Chd1 as indicated. As expected, *chd1*Δ and *rtf1*Δ strains have strong and modest cryptic initiation phenotypes, respectively (2). Strips within each vertical panel are from the same plate and organized for clarity (n=3). (**B**) Indirect immunofluorescence of untagged Chd1, HSV-Chd1, and HSV-Chd1ΔCHCT. Brightfield, DAPI, Alexa Fluor 568 (for HSV-Chd1), and merge of DAPI and Alexa Fluor 568 images of HSV antibody-treated cells are shown (n=3). (**C**) Left panels: Metaplots of average spike-in normalized RNA Pol II (8WG16), HA-Rtf1, and HSV-Chd1 ChIP-seq signal over the top 20% HSV-Chd1 occupied genes as determined from the wild-type dataset. Genes within this class with a maximum length of 1 kb after the +1 dyad are shown (n=324 genes). Dotted vertical lines show the maximum point of factor occupancy for each strain. Right panels: Violin plots to the right of corresponding metaplots demonstrate maximum point of factor occupancy for each gene; median and interquartile ranges for the data are shown. Wilcoxon-rank sum test was used to compare each strain to wild type: where * $p<0.05$, ** $p<0.01$, *** $p<0.001$, and **** $p<0.0001$. The average of 2-5 biological replicates for each strain is shown. Information on replicates can be found in the Supplementary Data File. (**D**) ChIP-qPCR analysis of HSV-Chd1 occupancy at the 5' and 3' ends of *CDC19, YEF3,* and *PMA1* in the indicated strains. ChIP-qPCR signals at the tested loci were normalized to those at *TELVI*. Unpaired t-tests comparing wild type to each mutant strain were performed. Asterisks represent p values where * $p<0.05$, ** $p<0.01$, and *** $p<0.001$. (**E**) ChIP-qPCR analysis of HA-Rtf1 occupancy as in D.

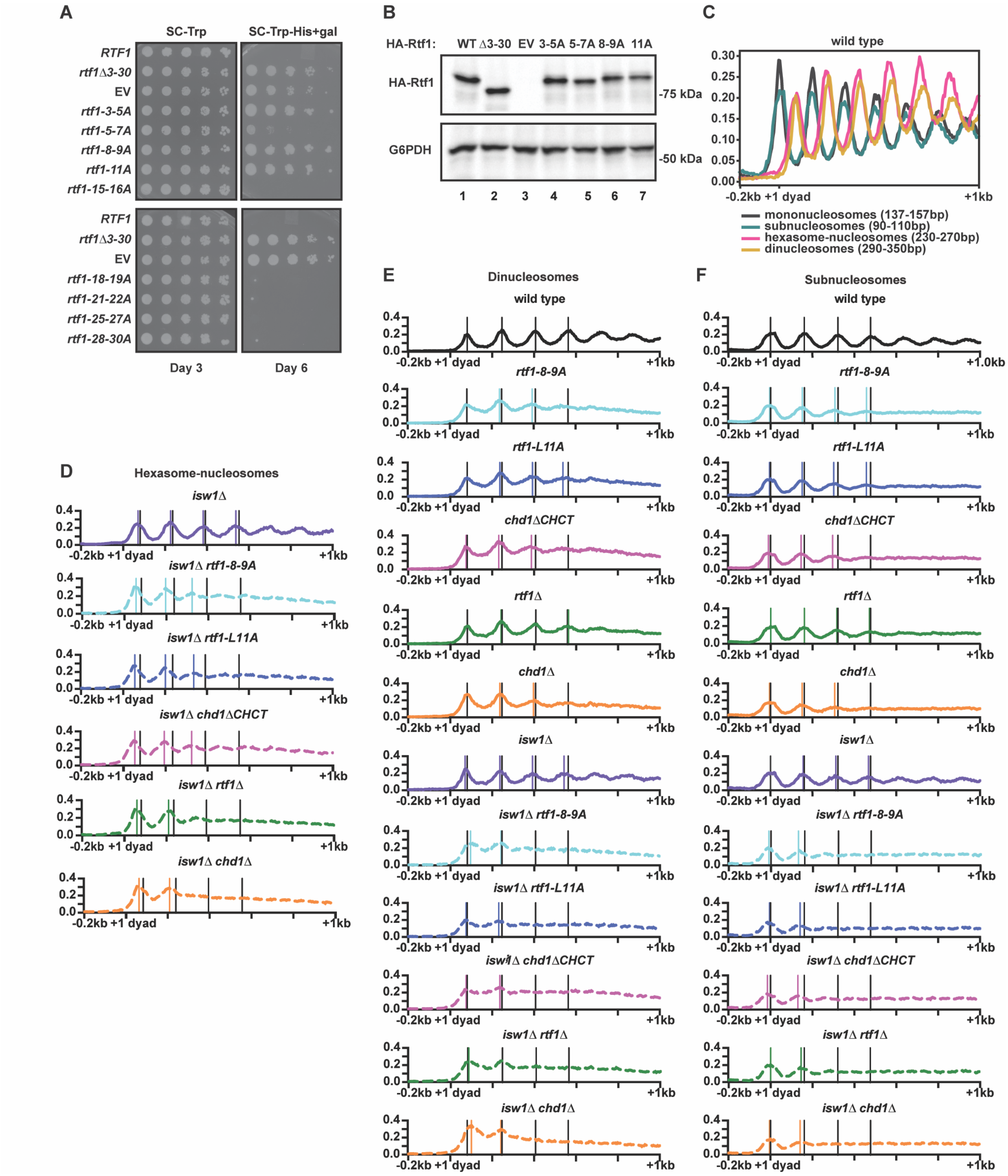


**Supplementary Figure S3. Substitutions in the Rtf1 LLALA box cause cryptic transcription initiation and alter chromatin structure over gene bodies.** (**A**) Analysis of cryptic transcription initiation using the *GAL1p-FLO8-HIS3* reporter in an *rtf1Δ* strain transformed with *CEN/ARS* plasmids expressing full-length HA-tagged Rtf1 protein with wild-type sequence, alanine substitutions, or a deletion of amino acids 3-30 (Δ3-30) (n=3). EV = empty vector control. (**B**) Western blot analysis of HA-Rtf1 wild-type and mutant proteins in the transformants tested for cryptic initiation in A. G6PDH served as the loading control. (**C**) Distribution of MNase-seq data over the top 20% HSV-Chd1 occupied genes spanning from 200bp upstream of the +1 nucleosome dyad to 1 kb downstream (maximum of 1 kb in length after +1 dyad, n=324 genes) in the wild-type strain. MNase-seq data were computationally size selected for mononucleosomes (137-157bp), subnucleosomes (90-110bp), hexasome-nucleosome complexes (230-270bp), and dinucleosomes (290-350bp). (**D-F**) Positions of hexasome-nucleosome complexes (**D**), dinucleosomes (**E**), and subnucleosome particles (**F**) as determined by MNase-seq analysis of chromatin isolated from the indicated yeast strains. Black and colored vertical lines denote the calculated center points in wild-type and mutant strains, respectively. Where colored lines are absent, the center point was either unchanged relative to wild type or unable to be calculated.

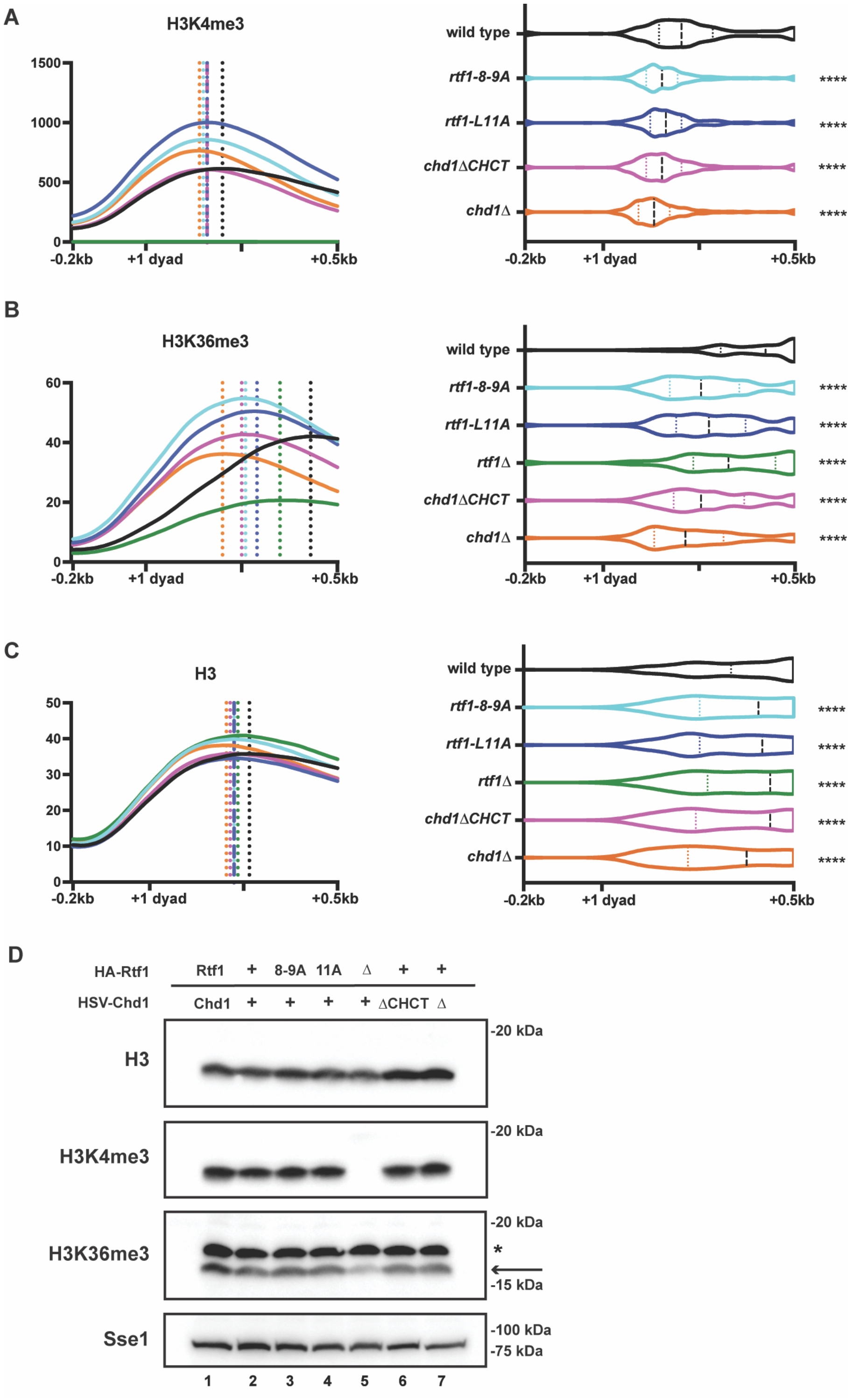


**Supplementary Figure S4. Levels and distributions of histone modifications in Rtf1-Chd1 interaction mutants.** (**A**) Metaplot of average spike-in normalized H3K4me3 ChIP-seq data over the top 20% of HSV-Chd1-occupied genes (determined from the wild-type dataset, under 1 kb in length after +1 dyad, n=324 genes) where dotted vertical lines indicate the maximum point of factor occupancy for each strain (left). Violin plots representing maximum point of factor occupancy for each individual gene and median and interquartile ranges for the data are shown (right). Wilcoxon-rank sum test was used to compare each strain to wild type: where * $p<0.05$, ** $p<0.01$, *** $p<0.001$, and **** $p<0.0001$. Data represent averages from 2-5 biological replicates. Information on replicates is provided in the Supplementary Data File. (**B**) As in A, for H3K36me3 ChIP-seq data. Data are averaged from 2-3 biological replicates. (**C**) As in A, for H3 ChIP-seq data. Data are averaged from 2-3 biological replicates. (**D**) Western blot analysis assessing levels of H3, H3K4me3, and H3K36me3 in wild-type and mutant strains. The relevant band in the H3K36me3 blot is indicated with an arrow and was determined by comparisons with a *set2*Δ strain. The slower migrating band, indicated with an asterisk, cross-reacts with the anti-H3K36me3 antibody. Sse1 served as the loading control.

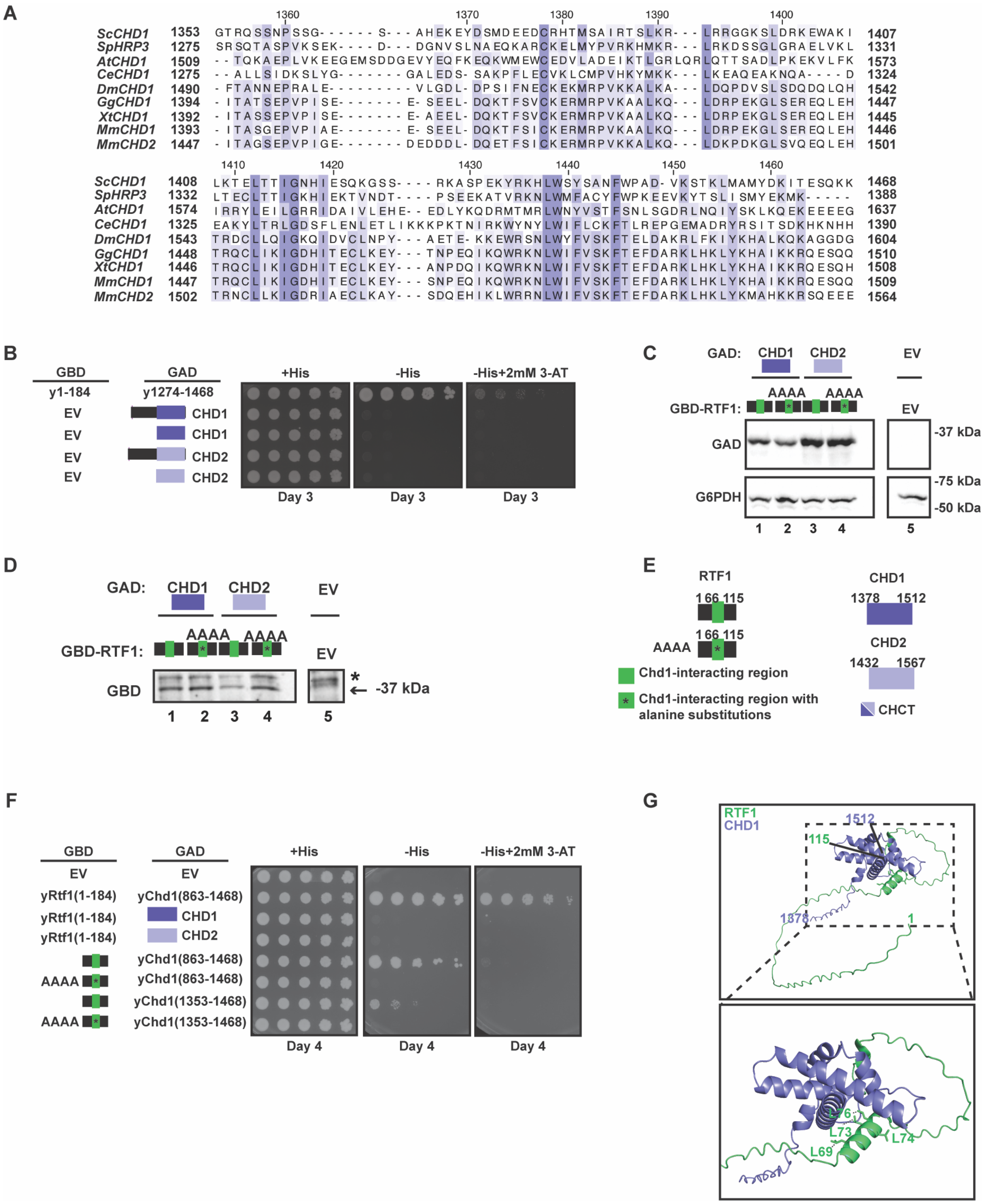


**Supplementary Figure S5. A cross-species interaction is observed between mouse RTF1 and the yeast Chd1 CHCT domain and is dependent on the RTF1 LLSLA box.** (**A**) Multiple sequence alignment of the *S. cerevisiae* Chd1 CHCT domain (amino acids 1353-1468) with Chd1 homologs from *S. pombe*, *A. thaliana*, *C. elegans*, *D. melanogaster*, *G. gallus*, *X. tropicalis*, and *M. musculus*. Also included is *M. musculus* CHD2. (**B**) Y2H analysis of GBD empty vector (EV) controls with CHD1 and CHD2 constructs defined and tested in Figure 5 (n=3). (**C**) Western blot analysis using anti-GAD antibody to assess levels of the GAD-CHD1 CHCT and GAD-CHD2 CHCT proteins in Y2H transformants containing either GBD-mRTF(1-115) or GBD-mRTF1(1-115, AAAA). The AAAA substitution in RTF1 is labeled above and indicated by an asterisk. G6PDH serves as a loading control. Cropped lanes are from the same membrane and are organized for clarity. (**D**) Western blot analysis using anti-GBD antibody to assess levels of the GBD-mRTF1(1-115) or GBD-mRTF1(1-115, AAAA) proteins in Y2H transformants containing either GAD-CHD1 CHCT or GAD-CHD2 CHCT. To increase the signal, whole cell extracts were first immunoprecipitated with a GBD antibody before resolving by SDS-PAGE. Cropped lanes are from the same membrane and are organized for clarity. Arrow indicates antibody-specific band and asterisk denotes background band. (**E**) Constructs used in the experiment shown in panel F to test for a Y2H interaction between yeast and mouse proteins. (**F**) Y2H analysis of interactions between yeast Rtf1 and mouse CHD1 or CHD2 and between mouse RTF1 and yeast Chd1. Plasmids expressing yeast GBD-Rtf1 and mouse GAD-CHD1 or CHD2 CHCT proteins and mouse GBD-RTF1 and yeast GAD-Chd1 proteins were co-transformed as indicated. The mouse proteins are represented graphically (see panel E). Cells were plated on media as indicated and images were taken after four days of growth (n=3). (**G**) AlphaFold 3 model predicting a potential interaction between mouse RTF1 (amino acids 1-115) and mouse CHD1 (amino acids 1378-1512) (3).

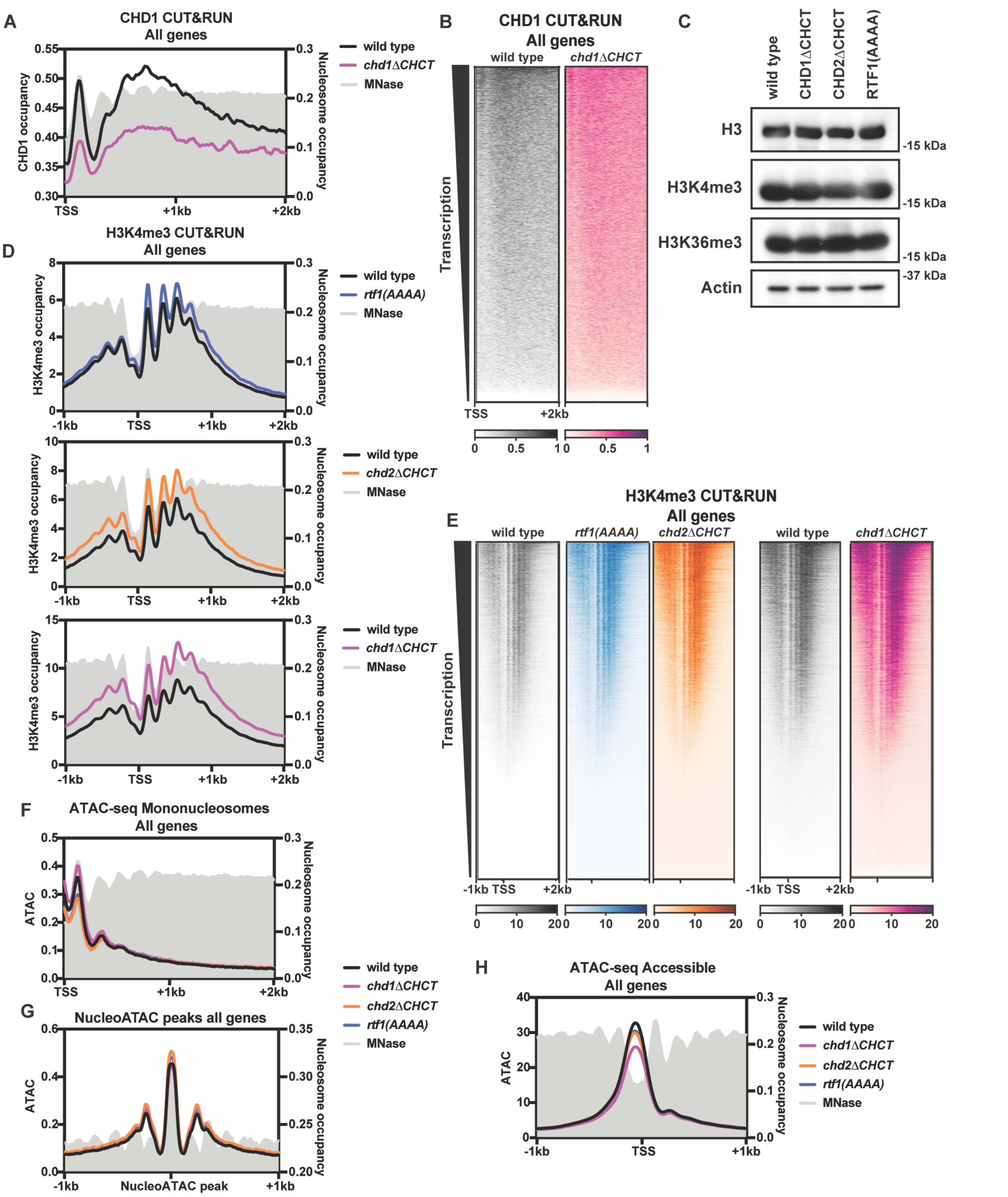


**Supplementary Figure S6. mES cell lines with mutations predicted to disrupt the mRTF1-mCHD1/2 interactions have minimal changes in H3K4me3 occupancy and chromatin accessibility.** (**A**) Metaplot of average CHD1 CUT&RUN data in wild-type (black) and *chd1ΔCHCT* (pink) cell lines over all genes. Data are averaged from three wild-type biological replicates and four *chd1ΔCHCT* replicates (two independently derived clones assayed in technical duplicate). The gray signal represents mononucleosome data from MNase-seq in wild-type mES cells (4). Left y-axis labels: relative CUT&RUN enrichment. Right y-axis labels: mononucleosome enrichment. (**B**) Heatmap of average CHD1 CUT&RUN data in wild-type (black) and *chd1∆CHCT* (pink) cell lines over all genes, sorted by transcription (5). Data are averaged from three wild-type biological replicates and four *chd1∆CHCT* replicates (as above). (**C**) Western blot analysis for H3, H3K4me3, and H3K36me3, in the indicated mES cell lines. Actin served as a loading control. (**D**) Metaplot of H3K4me3 CUT&RUN signal over all genes in wild-type (black), *rtf1*(*AAAA*) (blue), *chd1∆CHCT* (pink), and *chd2∆CHCT* (orange) cell lines. Y-axis labels as in A. Data are averaged from three wild-type, *rtf1*(*AAAA*), and *chd2∆CHCT* biological replicates and four *chd1∆CHCT* replicates (as above). Samples are paired with respective wild-type replicates for *rtf1*(*AAAA*) and *chd2∆CHCT* or *chd1∆CHCT*. (**E**) Heatmaps of H3K4me3 CUT&RUN signal over all genes sorted by transcription (5). (**F**) Metaplot of ATAC-seq mononucleosome (180-247bp) data over all genes . Left y-axis corresponds to ATAC-seq enrichment and right y-axis corresponds to mononucleosome enrichment as in A. Data are averaged from three wild-type, *rtf1*(*AAAA*), and *chd2∆CHCT* biological replicates and four *chd1∆CHCT* replicates (as above). (**G**) Mononucleosome (180-247 bp) ATAC-seq data over wild-type nucleosomes at all genes called using nucleoATAC (6). The gray signal represents mononucleosome data from MNase-seq in wild-type mES cells. (**H**) Metaplot of ATAC-seq accessible footprint (1-100bp) data at all genes as in F.

**Supplementary Table S1. *S. cerevisiae* and *S. pombe* strains used in this study**

| Alias | Mating type | Genotype |
|---|---|---|
| PJ69-4A (7) | a | *his3-200 leu2-3,112 ura3-52 trp1-901 gal4Δ gal80Δ LYS2::GAL1-HIS3 GAL2-ADE2 met2::GAL7-lacZ* |
| OKA309 | a | *his3-200 leu2-3,112 ura3-52 trp1-901 gal4Δ gal80Δ LYS2::GAL1-HIS3 GAL2-ADE2 met2::GAL7-lacZ chd1Δ* |
| KY1370 | α | *rtf1Δ::ARG4 KanMx::GAL1p-FLO8-HIS3 TEL-VR::URA3 his3Δ200 arg4-12 ura3-52 trp1Δ63* |
| KY2822 | a | *his3Δ200 lys2-128∂ leu2Δ1 ura3-52 trp1Δ63 KanMx::GAL1p-FLO8-HIS3 3xHSV-CHD1* |
| KY2937 | a | *his3Δ200 lys2-128∂ leu2Δ1 ura3-52 trp1Δ63 KanMx::GAL1p-FLO8-HIS3 3xHSV-CHD1 rtf1Δ::KanMx* |
| KY3804 | α | *his3Δ200 lys2-128∂ leu2Δ1 ura3-52 trp1Δ63 KanMx::GAL1p-FLO8-HIS3 3xHSV-CHD1 3xHA-RTF1* |
| KY3806 | a | *his3Δ200 lys2-128∂ leu2Δ1 ura3-52 trp1Δ63 3xHSV-CHD1 3xHA-RTF1* |
| KY3835 | α | *his3Δ200 lys2-128∂ leu2Δ1 ura3-52 trp1Δ63 KanMx::GAL1p-FLO8-HIS3* |
| KY3843 | α | *his3Δ200 lys2-128∂ leu2Δ1 ura3-52 trp1Δ63 KanMx::GAL1p-FLO8-HIS3 3xHSV-CHD1 3XHA-rtf1(8-9A)* |
| KY3844 | α | *his3Δ200 lys2-128∂ leu2Δ1 ura3-52 trp1Δ63 KanMx::GAL1p-FLO8-HIS3 3xHSV-CHD1 3XHA-rtf1(11A)* |
| KY3850 | α | *his3Δ200 lys2-128∂ leu2Δ1 ura3-52 trp1Δ63 KanMx::GAL1p-FLO8-HIS3 3xHSV-CHD1 rtf1Δ101::LEU2* |
| KY3853 | α | *his3Δ200 lys2-128∂ leu2Δ1 ura3-52 trp1Δ63 KanMx::GAL1p-FLO8-HIS3 chd1Δ::LEU2 3xHA-RTF1* |
| KY3934 | α | *his3Δ200 lys2-128∂ leu2Δ1 ura3-52 trp1Δ63 chd1Δ::LEU2 3xHA-RTF1 isw1Δ::HIS3* |
| KY3938 | α | *his3Δ200 lys2-128∂ leu2Δ1 ura3-52 3xHSV-CHD1 3XHA-rtf1(11A) isw1Δ::HIS3* |
| KY3950 | α | *his3Δ200 lys2-128∂ leu2Δ1 ura3-52 trp1Δ63 3xHSV-CHD1 3xHA-RTF1 isw1Δ::HIS3* |
| KY3956 | α | *his3Δ200 lys2-128∂ leu2Δ1 ura3-52 trp1Δ63 3xHSV-CHD1 3XHA-rtf1(8-9A) isw1Δ::HIS3* |

| | | |
|---|---|---|
| KY3984 | α | *his3Δ200 lys2-128∂ leu2Δ1 ura3-52 trp1Δ63 3xHSV-CHD1 rtf1Δ101::LEU2 isw1Δ::HIS3* |
| KY4361 | a | *his3Δ200 lys2-128∂ leu2Δ1 ura3-52 trp1Δ63 3xHSV-chd1(ΔCHCT)::pCORE-KU 3xHA-RTF1* |
| KY4363 | α | *his3Δ200 lys2-128∂ leu2Δ1 ura3-52 trp1Δ63 KanMx::GAL1p-FLO8-HIS3 3xHSV-chd1ΔCHCT 3xHA-RTF1* |
| KY4418 | α | *his3Δ200 lys2-128∂ leu2Δ1 ura3-52 trp1Δ63 3xHSV-chd1ΔCHCT 3xHA-RTF1 isw1Δ::HIS3* |
| KY5011 | α | *his3Δ200 lys2-128∂ leu2Δ1 ura3-52 trp1Δ63 KanMx::GAL1p-FLO8-HIS3 3xHSV-chd1ΔCHCT rtf1Δ::KanMx* |
| KY5466 | α | *his3Δ200 lys2-128∂ leu2Δ1 ura3-52 trp1Δ63 KanMx::GAL1p-FLO8-HIS3 3xHSV-CHD1 rtf1Δ::pCORE-UH* |
| FWP568 *(S. pombe)* | *h+* | *ura4-D18 leu1-32 ade6-m210 spt5+::3HA-KanMx* |
| KP08 *(S. pombe)* | *h+* | *ura4-D18 leu1-32 ade6-m210 spt5+::3HA-KanMx hrp1::HSV-hphMX6* |

**Supplementary Table S2. Oligonucleotide information for experiments conducted in *S. cerevisiae***

| Primer | Primer sequence (5' to 3') | Description |
|---|---|---|
| PCD01 | GACTGTATCGCCGGAATTCCCGGGGATCCCCATGTCTGCTGCAGCTGAGGATTTATTAGC | Assembly of GBD-Rtf1(1-30) D3A, L4A, D5A and HA-Rtf1(D3A, L4A, D5A) |
| PCD02 | GAAATTCGCCCGGAATTAGCTTGGCTG | Assembly of GBD-Rtf1(1-30) D3A, L4A, D5A; L8A, L9A; L11A; T25A, T26A, T27A; S28A, K30A; and HA-Rtf1(D3A, L4A, D5A) |
| MDH11 | GATCCCCATGTCTGATTTAGTTGCGGTTTTATTAGCCTTGGCTGGTG | Assembly of GBD-Rtf1(1-30) D5A, E6A, D7A |
| MDH12 | CACCAGCCAAGGCTAATAAAACCGCAACTAAATCAGACATGGGGATC | Assembly of GBD-Rtf1(1-30) D5A, E6A, D7A |
| SAT01 | GCTATTTCTACTGATTTTTCC | Assembly of GBD-Rtf1(1-30) L8A, L9A and GBD-Rtf1(1-30) L11A |
| SAT04 | ACCAGCCAAGGCGGCGGCATCCTCATCTAAATC | Assembly of GBD-Rtf1(1-30) L8A, L9A and HA-Rtf1(L8A, L9A) |
| SAT03 | GATTTAGATGAGGATGCCGCCGCCTTGGCTGGT | Assembly of GBD-Rtf1(1-30) L8A, L9A and HA-Rtf1(L8A, L9A) |
| SAT05 | GGATTTATTAGCCGCCGCTGGTGCCGATGAATCCG | Assembly of GBD-Rtf1(1-30) L11A and HA-Rtf1(L11A) |
| SAT06 | CGGCACCAGCGGCGGCTAATAAATCCTCATC | Assembly of GBD-Rtf1(1-30) L11A and HA-Rtf1(L11A) |
| MDH07 | GCCTTGGCTGGTGCCGCTGCATCCGAAGAAGAAGAT | Assembly of GBD-Rtf1(1-30) D15A, E16A and HA-Rtf1(D15A, E16A) |
| MDH08 | ATCTTCTTCTTCGGATGCAGCGGCACCAGCCAAGGC | Assembly of GBD-Rtf1(1-30) D15A, E16A and HA-Rtf1(D15A, E16A) |
| MDH15 | CTGGTGCCGATGAATCCGCAGCAGAAGATCAAGTTTTGAC | Assembly of GBD-Rtf1(1-30) E18A, E19A and HA-Rtf1(E18A, E19A) |
| MDH16 | GTCAAAACTTGATCTTCTGCTGCGGATTCATCGGCACCAG | Assembly of GBD-Rtf1(1-30) E18A, E19A and HA-Rtf1(E18A, E19A) |
| MDH17 | GCCGATGAATCCGAAGAAGAAGCTGCAGTTTTGACAACTACATCTGCC | Assembly of GBD-Rtf1(1-30) D21A, Q22A and HA-Rtf1(D21A, Q22A) |

| | | |
|---|---|---|
| MDH18 | GGCAGATGTAGTTGTCAAAACTG CAGCTTCTTCTTCGGATTCATCGG C | Assembly of GBD-Rtf1(1-30) D21A Q22A and HA-Rtf1(D21A Q22A) |
| PCD03 | GACAGTTGACTGTATCGCCGG | Assembly of GBD-Rtf1(1-30) T25A, T26A, T27A and GBD-Rtf1(1-30) S28A, K30A |
| PCD04 | GGCAGAAGCAGCAGCCAAAACTT GATCTTCTTCTTCGG | Assembly of GBD-Rtf1(1-30) T25A, T26A, T27A and HA-Rtf1(T25A, T26A, T27A) |
| PCD05 | GTTTTGGCTGCTGCTTCTGCCAAA TAGTAGGCAGG | Assembly of GBD-Rtf1(1-30) T25A, T26A, T27A and HA-Rtf1(T25A, T26A, T27A) |
| PCD06 | GCCTACTATGCGGCAGCTGTAGT TGTCAAAACTTG | Assembly of GBD-Rtf1(1-30) S28A, K30A |
| PCD07 | CAGCTGCCGCATAGTAGGCAGGA CAATTTCAGTG | Assembly of GBD-Rtf1(1-30) S28A, K30A |
| MDH13 | CTGCTCATATGTCTGATTTAGTTG CGGTTTTATTAGCCTTGGCTGGTG C | Assembly of HA-Rtf1(D5A, E6A, D7A) |
| MDH14 | GCACCAGCCAAGGCTAATAAAAC CGCAACTAAATCAGACATATGAGC AG | Assembly of HA-Rtf1(D5A, E6A, D7A) |
| SAT02 | CGAAGATTTCTTGACTCCTCAAAC AG | Assembly of HA-Rtf1(L8A, L9A) |
| PCD09 | CCTCTCCTGGTACTTCTGCATG | Assembly of HA-Rtf1(L8A, L9A), HA-Rtf1(L11A), HA-Rtf1(T25A, T26A, T27A), and HA-Rtf1(S28A, K30A) |
| SAT02 | CGAAGATTTCTTGACTCCTCAAAC AG | Assembly of HA-Rtf1(L11A) |
| PCD08 | GCACTAATTTGTTGAGAGCAC | Assembly of HA-Rtf1(T25A, T26A, T27A) and HA-Rtf1(S28A, K30A) |
| PCD14 | GCCCTTGCGGCAGCTGTAGTTGT CAAAACTTGATC | Assembly of HA-Rtf1(S28A, K30A) |
| PCD15 | CTACAGCTGCCGCAAGGGCAAAA AACAACGACC | Assembly of HA-Rtf1(S28A, K30A) |
| PSO551 | AGATCGAATTCCCGGGTACAAGG CAATCAAGCAACCC | Assembly of pGAD-Chd1(1353-1468) |
| HMO2 | CTCTGCAGGTCGACGTCACTTCTT TTGAGACTCTGT | Assembly of pGAD-Chd1(1353-1468) |

| | | |
|---|---|---|
| SAT60 | AAAGAGATCGAATTCCCGGGATCCCAAAGATTGTGTGGTGCTGGTGG | Assembly of GAD-mCHD1(1321-1506) |
| SAT61 | GATCTCTGCAGGTCGACGTCATTCTTGTCTCTTCTTGATAGCGTGCT | Assembly of GAD-mCHD1(1321-1506) |
| SAT62 | AAAGAGATCGAATTCCCGGGATCCGTTGCTTCTGGTGAAGAAGCTAAGTTG | Assembly of GAD-mCHD2(1331-1561) |
| SAT63 | GATCTCTGCAGGTCGACGTCATTGAGATCTCTTCTTGTGAGCCATCTTG | Assembly of GAD-mCHD2(1331-1561) |
| SAT73 | ATACCCCACCAAACCCAAAAAAAGAGATCGAATTCCCGGGGAAGGACAGATCTAAGAAGTCTGTTGTTTCT | Assembly of GAD-mCHD1(1378-1512) |
| SAT74 | TTTTCAGTATCTACGATTCATAGATCTCTGCAGGTCGACGTCAGTCAGAGTTTTGTTGAGATTCTTGTCTCTT | Assembly of GAD-mCHD1(1378-1512) |
| SAT75 | ATACCCCACCAAACCCAAAAAAAGAGATCGAATTCCCGGGGGGTAAGTCTTCTTCTAAGTCTAAGAGATCTCAAGG | Assembly of GAD-mCHD2(1432-1567) |
| SAT76 | TTTTCAGTATCTACGATTCATAGATCTCTGCAGGTCGACGTCACTTTTGTTCTTCTTCTTCTTGAGATCTCTTCTTGTGAGCCATCTTG | Assembly of GAD-mCHD2(1432-1567) |
| SAT68 | AAGGTCAAAGACAGTTGACTGTATCGCCGGAATTCCCGGGGATGAGAGGTAGATTGTGTGTTGGTAGAG | Assembly of GBD-mRTF1(1-321) and GBD-mRTF1(1-115) |
| SAT70 | TAAGAAATTCGCCCGGAATTAGCTTGGCTGCAGGTCGACGGATCCTCATTCGTCGTCAGAGTAAACTTCAGAAGT | Assembly of GBD-mRTF1(1-321) |
| SAT69 | TAAGAAATTCGCCCGGAATTAGCTTGGCTGCAGGTCGACGTCACTTGTTAGAACCGAAAGTCCATTCG | Assembly of GBD-mRTF1(1-115) |
| SAT77 | AGCGGCGGCTGCTGCTTCTTGGTCCAAGTTTTC | Assembly of GBD-mRTF1(1-115) L to A and GBD-mRTF1(1-321) L to A |
| SAT78 | GAAGCAGCAGCCGCCGCTAAGAGAAAGAGATCTGACTC | Assembly of GBD-mRTF1(1-115) L to A and GBD-mRTF1(1-321) L to A |

| | | |
|---|---|---|
| SAT64 | GACGAAAACTTGGACCAAGAATT GTTGTCTTTGGCTAAGAGAAAGA GATCTGACTCTGAAGAAAAGGAA CCACCAGTTTCTCAACCAGCTTGA | Assembly of GBD-mRTF1(66-95) |
| SAT65 | TCAAGCTGGTTGAGAAACTGGTG GTTCCTTTTCTTCAGAGTCAGATC TCTTTCTCTTAGCCAAAGACAACA ATTCTTGGTCCAAGTTTTCGTC | Assembly of GBD-mRTF1(66-95) |
| SAT66 | GCCGGAATTCCCGGGAGACGAAA ACTTGGACCAAGAATTGTTGT | Assembly of GBD-mRTF1(66-95) |
| SAT67 | GGCTGCAGGTCGACGGATCCTCA AGCTGGTTGAGAAACTGGTGG | Assembly of GBD-mRTF1(66-95) |
| SAT71 | AAGGTCAAAGACAGTTGACTGTAT CGCCGGAATTCCCGGGGGACGA AAACTTGGACCAAGAATTGTTG | Assembly of GBD-mRTF1(66-321) |
| SAT72 | TAAGAAATTCGCCCGGAATTAGCT TGGCTGCAGGTCGACGGATCCTC ATTCGTCGTCAGAGTAAACTTCAG AAGT | Assembly of GBD-mRTF1(66-321) |
| BTO64 | GCTTTCTGAACCAGTCCTG | To amplify *rtf1* locus for PCORE replacement cassettes |
| SAT09 | CTCATCCTTGTACTTTCCTTCCAA GGG | To amplify *rtf1* locus for PCORE replacement cassettes |
| SAT79 | GGTGCCATTCACTTGGGCAGAAG AG | To confirm pCORE-KU insertion; PCR to amplify from Chd1 gene through Chd1-1352aa |
| SAT80 | TCTCGATGAGCCCTTTTGCGATTC G | To confirm pCORE-KU insertion; PCR to amplify from Chd1 gene through Chd1-1352aa |
| SAT81 | TTGTTTCAATTATCTCTGGAAACC TTGGAC | PCR to amplify from Chd1 gene through Chd1-1352aa |
| SAT82 | GTCCAAGGTTTCCAGAGATAATTG AAACAATTTTTCTTCACCACATTTT CCATTGTTCCTTCCCCCC | PCR to amplify from stop to downstream genomic CHD1 region |
| SAT83 | CTGGAGCGAAAGAGAACAATTG | PCR to amplify from stop to downstream genomic CHD1 region |
| SAT84 | GACGAAAGAGGAAGATGAGAAG | Sequencing primers for final HSV-Chd1ΔCHCT |

| | | |
|---|---|---|
| SAT85 | GCTGTCTTATCAGTAATATCAGCCAT | Sequencing primers for final HSV-Chd1ΔCHCT |
| ECO238 | ACCAAGGGTCCAGAAATCAGAAC | qPCR location *CDC19* 5'; primer efficiency 1.98; (8) |
| ECO239 | TGTCATCGGTGGTGAAGATCAT | qPCR location *CDC19* 5'; primer efficiency 1.98; (8) |
| ECO240 | AGAAACTGTACTCCAAAGCCAACCT | qPCR location *CDC19* 3'; primer efficiency 1.99; (8) |
| ECO241 | TGGTCTGTACTTGGAAACCAATCTT | qPCR location *CDC19* 3'; primer efficiency 1.99; (8) |
| SAT35 | CCAGCAATCCATTAAGGTTC | qPCR location *YEF3* 5'; primer efficiency 2.01 |
| SAT36 | CAGCGGTCTTCTTGTCCTTG | qPCR location *YEF3* 5'; primer efficiency 2.01 |
| SAT31 | TTGGGTGCTTTGTCTAAGGC | qPCR location *YEF3* 3'; primer efficiency 2.02; |
| SAT32 | CGGCCCAGACTTCTTCAGTC | qPCR location *YEF3* 3'; primer efficiency 2.02; |
| ECO234 | GCTAGACCAGTTCCAGAAGAATATTTACA | qPCR location *PMA1* 5'; primer efficiency 1.97; (8) |
| ECO235 | CAGCCATTTGATTCAAACCGT | qPCR location *PMA1* 5'; primer efficiency 1.97; (8) |
| ECO236 | GAAATCTTCTTGGGTCTATGGATTG | qPCR location *PMA1* 3'; primer efficiency 1.94; (8) |
| ECO237 | CAACATCAGCGAAAATAGCGAT | qPCR location *PMA1* 3'; primer efficiency 1.94; (8) |
| APO95 | TGCAAGCGTAACAAAGCCATA | qPCR location *TELVI*; primer efficiency 2.0467; (8) |
| APO96 | TCCGAACGCTATTCCAGAAAG | qPCR location *TELVI*; primer efficiency 2.0467; (8) |

**Supplementary Table S3. Oligonucleotide information for CRISPR/Cas9 targeting in mES cells**

| Primer # | Primer sequence (5' to 3') | Primer purpose |
|---|---|---|
| 649 | CACCGaacctagcaacagtaaaggg | *CHD1* intron 30 cut F sgRNA cloning primer |
| 650 | AAACccctttactgttgctaggttC | *CHD1* intron 30 cut R sgRNA cloning primer |
| 651 | CACCGatccataagcctagatcagt | *CHD1* intron 33 cut F sgRNA cloning primer |
| 652 | AAACactgatctaggcttatggatC | *CHD1* intron 33 cut R sgRNA cloning primer |
| 761 | CACCGgtgtagaaaaaaagatcctg | *RTF1* LLSLA to AAAAA F sgRNA cloning primer |
| 762 | AAACcaggatctttttttctacacC | *RTF1* LLSLA to AAAAA R sgRNA cloning primer |
| 782 | CACCGtccctgctctagcaaagcac | *CHD2* intron 33 F sgRNA cloning primer |
| 810 | AAACgtgctttgctagagcagggaC | *CHD2* intron 33 R sgRNA cloning primer |
| 784 | CACCGgatttggtacagcctaagca | *CHD2* intron 36 F sgRNA cloning primer |
| 811 | AAACtgcttaggctgtaccaaatcC | *CHD2* intron 36 R sgRNA cloning primer |
| 655 | GGAGGAATTGGTTTGAGCACTCT | *CHD1ΔCHCT* PCR genotyping forward primer |
| 656 | GTGAGAACATCTTTGCAAGTG | *CHD1ΔCHCT* PCR genotyping reverse primer |
| 657 | GACACCATCACAGCTGGAACATG | *CHD1ΔCHCT* PCR genotyping reverse primer |
| 658 | GTTTGCATGGGAATATTGGTGATCTG | *RTF1* LLSLA sequence PCR genotyping forward prime |
| 659 | GATTTAGACAACAACAGCTTCC | *RTF1* LLSLA sequence PCR genotyping forward prime |
| 839 | CCAGACACACCTCCTCCATGCTTTT | *CHD2ΔCHCT* PCR genotyping forward primer |
| 840 | CCCGATGGGAACAGGCTCACT | *CHD2ΔCHCT* PCR genotyping reverse primer |
| 841 | GGGCCCACACAAGAATGGGACT | *CHD2ΔCHCT* PCR genotyping reverse primer |

**Supplementary Table S4. Recombinant DNA purchased for codon-optimized mouse Y2H experiments and for homology constructs in CRISPR/Cas9 targeting**

| **Purpose** | **Codon-optimized sequences for Y2H plasmids** |
|---|---|
| Codon optimized sequence corresponding to mouse CHD1(1321-1506) | CAAAGATTGTGTGGTGCTGGTGGTTCTAAGAGAAGAAAGACTAGAGC<br>TAAGAAGTCTAAGGCTATGAAGTCTATCAAGGTTAAGGAAGAAATCAA<br>GTCTGACTCTTCTCCATTGCCATCTGAAAAGTCTGACGAAGACGACGA<br>CAAGTTGAACGACTCTAAGCCAGAATCTAAGGACAGATCTAAGAAGTC<br>TGTTGTTTCTGACGCTCCAGTTCACATCACTGCTTCTGGTGAACCAGT<br>TCCAATCGCTGAAGAATCTGAAGAATTGGACCAAAAGACTTTCTCTAT<br>CTGTAAGGAAAGAATGAGACCAGTTAAGGCTGCTTTGAAGCAATTGG<br>ACAGACCAGAAAAGGGTTTGTCTGAAAGAGAACAATTGGAACACACT<br>AGACAATGTTTGATCAAGATCGGTGACCACATCACTGAATGTTTGAAG<br>GAATACTCTAACCCAGAACAAATCAAGCAATGGAGAAAGAACTTGTGG<br>ATCTTCGTTTCTAAGTTCACTGAATTCGACGCTAGAAAGTTGCACAAG<br>TTGTACAAGCACGCTATCAAGAAGAGACAAGAATGA |
| Codon optimized sequence corresponding to mouse CHD2(1331-1561) | GTTGCTTCTGGTGAAGAAGCTAAGTTGAAGAAGAGAAAGCCAAGAGT<br>TAAGAAGGAAAACAAGGCTCCAAGATTGAAGGACGAACACGGTTTGG<br>AACCAGCTTCTCCAAGACACTCTGACAACCCATCTGAAGAAGGTGAA<br>GTTAAGGACGACGGTTTGGAAAAGTCTCCAACTAAGAAGAAGCAAAA<br>GAAGAAGGAAAACAAGGAAAACAAGGAAAAGCCAGTTTCTTCTAGAA<br>AGGACAGAGAAGGTGACAAGGAAAGAAAGAAGTCTAAGGACAAGAAG<br>GAAAAGGTTAAGGGTGGTGACGGTAAGTCTTCTTCTAAGTCTAAGAG<br>ATCTCAAGGTCCAGTTCACATCACTGCTGGTTCTGAACCAGTTCCAAT<br>CGGTGAAGACGAAGACGACGACTTGGACCAAGAAACTTTCTCTATCT<br>GTAAGGAAAGAATGAGACCAGTTAAGAAGGCTTTGAAGCAATTGGAC<br>AAGCCAGACAAGGGTTTGTCTGTTCAAGAACAATTGGAACACACTAGA<br>AACTGTTTGTTGAAGATCGGTGACAGAATCGCTGAATGTTTGAAGGCT<br>TACTCTGACCAAGAACACATCAAGTTGTGGAGAAGAAACTTGTGGATC<br>TTCGTTTCTAAGTTCACTGAATTCGACGCTAGAAAGTTGCACAAGTTG<br>TACAAGATGGCTCACAAGAAGAGATCTCAATGA |
| Codon optimized sequence corresponding to mouse RTF1(1-321) | ATGAGAGGTAGATTGTGTGTTGGTAGAGCTGCTGCTGTTGCTGCTGC<br>TGTTGCTGCTGCTGCTGTTGCTGTTCCATTGGCTGGTGGTCAAGAAG<br>GTTCTCAAGGTGGTGTTAGAAGAGGTTCTAGAGGTACTACTATGGTTA<br>AGAAGAGAAAGGGTAGAGTTGTTATCGACTCTGACACTGAAGACTCT<br>GGTTCTGACGAAAACTTGGACCAAGAATTGTTGTCTTTGGCTAAGAGA<br>AAGAGATCTGACTCTGAAGAAAAGGAACCACCAGTTTCTCAACCAGCT<br>GCTTCTTCTGACTCTGAAACTTCTGACTCTGACGACGAATGGACTTTC<br>GGTTCTAACAAGAACAAGAAGAAGGGTAAGACTAGAAAGGTTGAAAA<br>GAAGGGTGCTATGAAGAAGCAAGCTAACAAGGCTGCTTCTTCTGGTT<br>CTTCTGACAGAGACTCTTCTGCTGAATCTTCTGCTCCAGAAGAAGGTG<br>AAGTTTCTGACTCTGAATCTTCTTCTTCTTCTTCTTCTGACTCTGA<br>CTCTTCTTCTGAAGACGAAGAATTCCACGACGGTTACGGTGAAGACTT<br>GATGGGTGACGAAGAAGACAGAGCTAGATTGGAACAAATGACTGAAA<br>AGGAAAGAGAACAAGAATTGTTCAACAGAATCGAAAAGAGAGAAGTTT<br>TGAAGAGAAGATTCGAAATCAAGAAGAAGTTGAAGACTGCTAAGAAG<br>AAGGAAAAGAAGGAAAAGAAGAAGAAGCAAGAAGAAGAACAAGAAAA<br>GAAGAAGTTGACTCAAATCCAAGAATCTCAAGTTACTTCTCACAACAA |

| | |
|---|---|
| | GGAAAGAAGATCTAAGAGAGACGAAAAGTTGGACAAGAAGTCTCAAG<br>CTATGGAAGAATTGAAGGCTGAAAGAGAAAAGAGAAAGAACAGAACT<br>GCTGAATTGTTGGCTAAGAAGCAACCATTGAAGACTTCTGAAGTTTAC<br>TCTGACGACGAATGA |
| **Purpose** | **Homology construct sequence** |
| *chd1∆CHCT* homology construct | atgtaggtagaacattgtatatatagtaaataaacaaatcttgtaaggaaaagaaaaaacaaaagaa<br>gctatgcagaaagaaaaagatggaagggaaggctggtattctagtgtaaaacctagaaagtggaaa<br>agcccaaactagtttcagtggctgagttctttagaagttgttgtgggcatataattgaggggaggaattggt<br>ttgagcactcttttaataccttacaagataatattacaaagcagacattctttgtcattgtttagcaaagtagc<br>aaggaaattttatttaaaaacctagcaacagtaaagggAAgtcatgtttgagggtcaagagagatctct<br>gagggaagttgggagaaaatgaagtatggctgttaacggagaataagtacttgcaaaataaaacatg<br>agtggttgttgtaaagagcatcacgttgtgtttatggttgggcttttgtttctttttcccagctgaatgactccaa<br>gcctgaaagtgtaattggtatataatattttattctttaaattatatcattaacaccatacatattctttattattta<br>gaaataaatggagaatttagttaataaaatagaggtaatgaataaattaaatcaaataccattggagtta<br>atcccataaaataatagtccaaagcataaatccataagcctagatcagtAgAcaaggaagcaggat<br>attatattggtctcatggataataattataaaaggagcaggatattcacatgaaaagagaaactgtttga<br>cattaccttaagtaatcacaacaaaccagaacatgaagacgtcggctacagtgcagagaaacatgtt<br>ccagctgtgatggtgtcaccaatagtataggatttgaatctagccatgggaacatttcagacaaactgca<br>ttagagggattctcaataaaagtactgaataagactcataaaaatgagccaggcgtggtggcgcatgc<br>ctttaatcccagcactcggga |
| *rtf1*(AAAA) replacement homology construct | gtgtgtttgatgtgtgcatatgggctgaaatacatgtggaagtctgaggacagcttcatgaagtcttttctct<br>ccttccactttttcgtggattctgaggaccaaactccagtcatcaggcttttccagcaagcactgagccgtc<br>tcactagccctctagatgcttccttatctttaaatgaggatgatgacaatatatattgtgggttgttgtgaaga<br>gttaagtgtcagatgctgagagacttagctggcatctagaaagaacagtaattgtcggctggtattgttatt<br>gcagggagtggaaccagtgaccctttgtcatgtcttttaataactggactgagtatatgggagaagtagt<br>gggaaaccatgagacagcagacaaccaccatcatcaccactaccaagacaaggtagagatggtg<br>aatgggagtgttgggatttttggattagaattctggccagcatcctccttactttgtgactttggactatatcctt<br>cagaaatataactcctttttctattttacattttcaaaatataaaatgacctcagaaacttatagtttgtaagct<br>taggtagaatagcttaggtagctacaggtgctaattttactatggaagacaataaaagatccttggagtgt<br>tgactatctgccatttaaaacatgaagcagattccctttgcatccatagctctggctgttcagtatgggtggt<br>cagtacttagcagtttagagagaaatctgaaagcttcaagtagctccagttccatttcctatttcactccctt<br>aaaggaggacagattttggagtttaagtttgcatgggaatattggtgatctgtctctgctcttgtgctcagtg<br>cttatgggtgtgtacttcaggtgaaagcacaggcgtgtagaaaaaaagatcctggAAagaatggcca<br>tttcattacacttttctggtgagcaaaactgacgtgaccatttctgcccaattaggagGCCGCTGCC<br>GCTgcgaaacggaagcgcagtgactccgaggagaaggagcctcctgtgagtcagcctgcagcct<br>catcagattcagagacctccgacagtgatgatgaggtgagtgtggaggagttaggcctggatgattgg<br>cgagtgtttgtggaagggtcctgacttggtgatgctgccttaaaccaagactgtaacctaagacctgaca<br>gagctgggtagggaacagaaattgtgtgagcagacctttacattatggtggccattctctgttttagaagg<br>gccaagcgttagtctttttcttttttttaaagatttatttatttatgagtgctttatctgtatgtacacttgcatgcca<br>gaagagggcatcagatcctagtatagatagttgtgagccaccatgtggttcctgggaattaaactcaga<br>acctctagaagaacagatagtgctcttaaccactgagctgtctctttaggcctccccccacccctccccc<br>catcattgttttaaaaagtcatcttttaaatactgcttttatgttagaaaacaaattctaagtcaggcattgtga<br>tgcatgcctataatttcagcattcaggaagctgaaacagtgggatcaagagttcaaagccagcctgagt<br>tatataggagaaaaattatacataaaaaatgtatatgctatatttattttagtgataaatactttgagacaca<br>aagcttctttggggagattgatatgccattcctgaaagtaatggtattgactttagaggaaggtaaaaag<br>gaagctgttgttgtctaaatcaaggcagttttctcaaattgtcca |

| | |
|---|---|
| *chd2∆CHCT* homology construct | tgtcagattccctgggcatttctgagcctccatgaaggtcctggaaaccatgccaggatcctctttgtaag<br>atcaagcattgttaactctaggactatctttccagacacacctcctccatgctttttaaaatttgtatttaatct<br>gtatcaaattttcctaaataatctgtgtgtgtccctctgaaggctttgaatctacattattttggtctccatgcttg<br>tttctcttggcttctgtcagttacagttaatatgcattcctggaccagactgtagctgttaataaagcttgctgc<br>atggtaattaggagttctgtttaaatccttagtttcttcatgaaatttctgtaaattatattgataaactcaaagc<br>tgtagttgagtacctttagacagaaatgggaatacttagaatttaggtattaatcttttaagtttgtgtgaattg<br>aggtcatcatttccctgctctagcaaagcactAAtcttgcagaggacccaggttcagttcgcagcaccc<br>acccacgttctgattcacagtgtttatgactccagtttctggagatctggcaccctctatggccttttcatgttc<br>atgcacagatgtggagcacataaacccatacagatacacacacagtcatataaaacagtaccacca<br>ccaccgtcaccaaataaaacaaacaaacagaactctgcctagattttaagattatttagtgatcatgact<br>ctttgtaactctttgaaaaggtaaaaacgcaaatgtagtcctaatttatacaactattgcaagtcagttcca<br>caggaaaccccttaggttttgtttttctctggggaggtaatgatcaatccgtggggttgttggtgccctatag<br>gccattgactctttgcctgtgttttacatggatttggtacagcctaagcaaAAtgacaaaaaccagcgca<br>ggtcacagagtgtgtgacctatggtccagacataatgtcatagactgtgatttttcattatactaggtaattt<br>cattagttcctaacatactcccatgagtaggcagacagtaaaacttttgtttattaggtaagataaccaatg<br>cattagagccaaatagggttttaaatcctgacaaggatctgttaacctgtgatctcatgacctggatgcttg<br>ttttctagacccttccttcttatgtatatcctttttttttagcttgtaatgtggcttgtcataagtcccattcttgtgtgg<br>gcccatgtacttggcatccatccagctcctcacacagaagattccaaattgctcacttaaaggagtaga<br>cagtttgcagtgcccagtagttgttgcccttttgtataagtacgtatgtattgtatggggatggccataaggg<br>tgacttggcttctgctgtagcattacttctcacgcaggcaagctctcctgtagatctgctaccgtcccacatg<br>ggccatgtccaaatcagccactgaggagcaattgtggacatt |

**Supplementary Table S5. Plasmids used for CRISPR/Cas9 targeting**

| **Purpose** | **Parent plasmid** | **Bacterial Resistance Cassette** | **Mammalian Resistance Cassette** |
|---|---|---|---|
| sgRNA for *CHD1* exon 30 | gRNA CRISPR vector pX330-U6-Chimeric_BB-CBh-hSpCas9 | $Amp^R$ | puromycin |
| sgRNA for *CHD1* exon 33 | gRNA CRISPR vector pX330-U6-Chimeric_BB-CBh-hSpCas9 | $Amp^R$ | puromycin |
| *chd1ΔCHCT* homology construct | TOPO vector | $Amp^R$ and $Kan^R$ | none |
| sgRNA for *RTF1* LLSLA for AAAAA | gRNA CRISPR vector pX330-U6-Chimeric_BB-CBh-hSpCas9 | $Amp^R$ | puromycin |
| *RTF1*(AAAA) homology construct | pMK-RQ GeneArt | $Kan^R$ | none |
| sgRNA for *CHD2* intron 33 | gRNA CRISPR vector pX330-U6-Chimeric_BB-CBh-hSpCas9 | $Amp^R$ | puromycin |
| sgRNA for *CHD2* intron 36 | gRNA CRISPR vector pX330-U6-Chimeric_BB-CBh-hSpCas9 | $Amp^R$ | puromycin |
| *chd2ΔCHCT* homology construct | TOPO vector | $Amp^R$ | none |

**Supplementary Table S6. Bacterial and yeast expression plasmids for experiments conducted in *S. cerevisiae***

| Plasmid | Insert description | *E. coli* marker | *S. cerevisiae* marker | Type | Purpose |
|---|---|---|---|---|---|
| pGBT9 (9) | GBD-EV | $Amp^R$ | *TRP1* | 2µ | Yeast two-hybrid |
| KB1271/ pARL01 | GBD-Rtf1(1-30) | $Amp^R$ | *TRP1* | 2µ | Yeast two-hybrid |
| KB1279/ pARL03 | GBD-Rtf1(1-184) | $Amp^R$ | *TRP1* | 2µ | Yeast two-hybrid |
| KB1552 | GBD-Rtf1(1-30) D3A, L4A, D5A | $Amp^R$ | *TRP1* | 2µ | Yeast two-hybrid |
| KB1375 | GBD-Rtf1(1-30) D5A, E6A, D7A | $Amp^R$ | *TRP1* | 2µ | Yeast two-hybrid |
| KB1655 | GBD-Rtf1(1-30) L8A, L9A | $Amp^R$ | *TRP1* | 2µ | Yeast two-hybrid |
| KB1656 | GBD-Rtf1(1-30) L11A | $Amp^R$ | *TRP1* | 2µ | Yeast two-hybrid |
| KB1373 | GBD-Rtf1(1-30) D15A, E16A | $Amp^R$ | *TRP1* | 2µ | Yeast two-hybrid |
| KB1376 | GBD-Rtf1(1-30) E18A, E19A | $Amp^R$ | *TRP1* | 2µ | Yeast two-hybrid |
| KB1377 | GBD-Rtf1(1-30) D21A, Q22A | $Amp^R$ | *TRP1* | 2µ | Yeast two-hybrid |
| KB1564 | GBD-Rtf1(1-30) T25A, T26A, T27A | $Amp^R$ | *TRP1* | 2µ | Yeast two-hybrid |
| KB1565 | GBD-Rtf1(1-30) S28A, K30A | $Amp^R$ | *TRP1* | 2µ | Yeast two-hybrid |
| pGAD424 (9) | GAD-EV | $Amp^R$ | *LEU2* | 2µ | Yeast two-hybrid |
| KB329 (1) | GAD-Chd1(863-1468) | $Amp^R$ | *LEU2* | 2µ | Yeast two-hybrid |
| KB1281 | GAD-Chd1(1274-1468) | $Amp^R$ | *LEU2* | 2µ | Yeast two-hybrid |

| | | | | | |
|---|---|---|---|---|---|
| KB1757 | GAD-Chd1(1353-1468) | $Amp^R$ | *LEU2* | 2µ | Yeast two-hybrid |
| KB1700 | GAD-mCHD1(1321-1506) | $Amp^R$ | *LEU2* | 2µ | Yeast two-hybrid |
| KB1701 | GAD-mCHD2(1331-1561) | $Amp^R$ | *LEU2* | 2µ | Yeast two-hybrid |
| KB1704 | GAD-mCHD1(1378-1512) | $Amp^R$ | *LEU2* | 2µ | Yeast two-hybrid |
| KB1705 | GAD-mCHD2(1432-1567) | $Amp^R$ | *LEU2* | 2µ | Yeast two-hybrid |
| KB1718 | GBD-mRTF1(1-321) | $Amp^R$ | *TRP1* | 2µ | Yeast two-hybrid |
| KB1719 | GBD-mRTF1(1-115) | $Amp^R$ | *TRP1* | 2µ | Yeast two-hybrid |
| KB1724 | GBD-mRTF1(1-115) L to A | $Amp^R$ | *TRP1* | 2µ | Yeast two-hybrid |
| KB1725 | GBD-mRTF1(1-321) L to A | $Amp^R$ | *TRP1* | 2µ | Yeast two-hybrid |
| KB1717 | GBD-mRTF1(66-95) | $Amp^R$ | *TRP1* | 2µ | Yeast two-hybrid |
| KB1720 | GBD-mRTF1(66-321) | $Amp^R$ | *TRP1* | 2µ | Yeast two-hybrid |
| KB1095/ pLH157-*LEU2* (10) | *E. coli* Tyr tRNA synthetase; *E. coli* tRNATyr amber suppressor tRNA | $Amp^R$ | *LEU2* | 2µ | BPA crosslinking |
| KB851/ pAP45 (8) | c-Myc-Rtf1 | $Kan^R$ | *TRP1* | 2µ | BPA crosslinking |
| KB1424 | c-Myc-Rtf1(L4 amber codon) | $Kan^R$ | *TRP1* | 2µ | BPA crosslinking |
| pRS314 (11) | pBluescript derivative, EV | $Amp^R$ | *TRP1* | *CEN/ARS* | Cryptic initiation test |
| KB660/ pLS21-5 (12) | 3xHA-RTF1 | $Amp^R$ | *TRP1* | *CEN/ARS* | Cryptic initiation test |

| | | | | | |
|---|---|---|---|---|---|
| KB661 (13) | 3xHA-Rtf1(Δ3-30) (i.e. *3xHA-rtf1Δ1*) | $Amp^R$ | *TRP1* | *CEN/ARS* | Cryptic initiation test |
| KB1553 | 3xHA-Rtf1(D3A, L4A, D5A) | $Amp^R$ | *TRP1* | *CEN/ARS* | Cryptic initiation test |
| KB1379 | 3xHA-Rtf1(D5A, E6A, D7A) | $Amp^R$ | *TRP1* | *CEN/ARS* | Cryptic initiation test |
| KB1653 | 3xHA-Rtf1(L8A, L9A) | $Amp^R$ | *TRP1* | *CEN/ARS* | Cryptic initiation test |
| KB1654 | 3xHA-Rtf1(L11A) | $Amp^R$ | *TRP1* | *CEN/ARS* | Cryptic initiation test |
| KB1555 | 3xHA-Rtf1(D15A, E16A) | $Amp^R$ | *TRP1* | *CEN/ARS* | Cryptic initiation test |
| KB1364 | 3xHA-Rtf1(E18A, E19A) | $Amp^R$ | *TRP1* | *CEN/ARS* | Cryptic initiation test |
| KB1380 | 3xHA-Rtf1(D21A, Q22A) | $Amp^R$ | *TRP1* | *CEN/ARS* | Cryptic initiation test |
| KB1566 | 3xHA-Rtf1(T25A, T26A, T27A) | $Amp^R$ | *TRP1* | *CEN/ARS* | Cryptic initiation test |
| KB1567 | 3xHA-Rtf1(S28A, K30A) | $Amp^R$ | *TRP1* | *CEN/ARS* | Cryptic initiation test |
| KB1575 (14) | pCORE (*K. lactis URA3*, *kanMX4*) | $Amp^R$ | | Bacterial expression | |
| KB1577 (14) | pCORE-UH *(K. lactis URA3*, *hphMX6)* | $Amp^R$ | | Bacterial expression | Site-specific mutagenesis at *RTF1* |
